# Profiling peripheral immune cells in Parkinson’s disease: A Scoping Review

**DOI:** 10.64898/2026.02.17.706426

**Authors:** Sherilyn Junelle Recinto, Janna E. Jernigan Posey, Nadejda Lefter, Zagorka Vitic, Morten Øgelund Overgaard, Lin Liu, Andrew J.M. Howden, Klaus Eyer, Marina Romero-Ramos, Malú Gámez Tansey, Jo Anne Stratton

## Abstract

Parkinson’s disease (PD) is increasingly recognized as a multi-system disorder with immune dysregulation extending beyond the central nervous system. Although numerous studies have examined peripheral immune alterations in people with PD, findings remain heterogenous and difficult to reconcile. To clarify the current landscape, we conducted a comprehensive scoping review of human studies profiling peripheral blood immune cells in PD. Following PRISMA-ScR guidelines, we systematically screened the literature and curated studies reporting *in vivo* and *ex vivo* immune characterizations from PD patients. Eligible studies based on pre-defined criteria were assessed for patient demographics and clinical variables, experimental and analytical approaches, and reported immune outcomes. Our synthesis reveals a steady expansion and diversification of peripheral immune cell research in PD especially over the last decade. Deep immunophenotyping identifies convergent signatures across *in vivo* studies of both innate and adaptive compartments, including expanded pro-inflammatory T-cell subsets, altered monocyte subset distributions, increased cytotoxic natural killer cells and neutrophil-to-lymphocyte ratio, and dysregulated pathways related to immune activation, chemotaxis, mitochondrial function, and autophagy-lysosomal processes. Stimulation-based *ex vivo* assays further demonstrate recurrent T-cell hyper-responsiveness in PD, whereas myeloid cell responses are more variable and context dependent. Critically, this review highlights substantial variability and under-reporting in study design, which impeded our ability to make strong conclusions relating to many aspects of PD peripheral immunity.

## Introduction

As the second most common and fastest growing neurodegenerative disorder, Parkinson’s Disease (PD) represents a major public health concern (Dorsey et al., 2018; Gadhave et al., 2024). It is projected to impact 25.2 million people worldwide by 2050, representing a 112% increase from 2021 (Su et al., 2025). People with PD (PwPD) experience a wide range of symptomology including sensory defects, affective disorders, cognitive decline or dementia, sleep disturbances, autonomic dysfunction that can impact the cardiovascular, gastrointestinal, genitourinary, and thermoregulatory systems, and characteristic motor impairments such as tremor, bradykinesia, rigidity, and postural instability that worsen with disease duration (Kalia & Lang, 2015). Treatment options so far are limited to symptomatic alleviation and do not halt or reverse the progression of the disease.

Historically, PD research has focused on neuropathological changes within the central nervous system (CNS), however, PD is now recognized as a multi-system disorder involving both central and peripheral immune alterations. A growing body of work implicates peripheral inflammation, gut-brain axis perturbations, and immune cell infiltration in disease initiation and progression, with Braak staging supporting a model in which pathology may spread from the gastrointestinal tract to the brain. Immune dysregulation has been documented across clinical, imaging, and molecular studies, including microglial dysfunction (Imamura et al., 2003; McGeer et al., 1988), elevated cytokines and chemokines (Delgado-Alvarado et al., 2017; Lindqvist et al., 2013), blood-brain-barrier disruption (Gray & Woulfe, 2015; Ham et al., 2014), and inflammatory changes in blood and cerebrospinal fluid (CSF) (Hallqvist et al., 2024; Mollenhauer et al., 2019; Qu et al., 2023). Moreover, circulating immune cell analyses indicate that inflammatory activity may peak early in disease to decrease or fail at later stages (Capelle et al., 2023; Farmen et al., 2021; Johansson et al., 2025; Konstantin Nissen et al., 2022; Lindestam Arlehamn et al., 2020; Mark, Titus, et al., 2025; Yan et al., 2021; Zhang et al., 2025). Collectively, these findings suggest that immune alterations are not merely secondary to neurodegeneration but may play causal roles in PD pathogenesis.

Despite this convergence, the PD immune literature remains fragmented and, at times, inconsistent. While changes in the amounts of cytokines/chemokines in biofluids may reflect the participation of peripheral immunity in disease mechanisms, their interpretation is complicated by pre-analytical and biological variability, including assay sensitivity and environmental factors, such as infections, circadian rhythms and diet (D’Esposito et al., 2022; Marchewka et al., 2024; Nguyen et al., 2021; Nilsonne et al., 2016). Furthermore, these measurements alone provide limited insight into the cellular sources and underlying molecular pathways driving immune alterations in PD. To overcome these challenges, it is crucial to directly assess changes in immune cell populations – their frequencies, phenotypes, and activation states. Such studies offer greater resolution for elucidating specific immune cell subsets involved, and the usage of functional stimulation-based *ex vivo* assays may uncover immune phenotypes that could otherwise be challenging to decipher at baseline and could be affected by exogenous factors.

That said, as with the study of any complex immunological disease, immune cell profiles and inferred mechanisms vary across studies, likely due to both biological and technical causes. Disease heterogeneity, patient demographics, disease stages/subtypes, differences in assay platforms, sample handling, analytical methods, and more, contribute to challenges in drawing definitive conclusions relating alterations in peripheral immunity in PD. Here we aimed to perform a comprehensive scoping review as an essential first step toward identifying common patterns, methodological gaps, and opportunities for standardization within the PD peripheral immune cell literature. We conducted a comprehensive search using the PubMed database by employing keywords and Medical Subject Headings (MeSH) including those related to “Parkinson’s disease”, “periphery” and “inflammation”. Where some studies were not identified based on this search criteria, additional studies were included based on expert reviewer knowledge. Titles, abstracts and full texts were screened, and data were extracted by multiple independent researchers using a curated set of criteria such as patient demographics, experimental design and study rigor. We highlight reproducible immune features across studies, while emphasizing the need for standardized reporting practices and cell-based analytical approaches to advance our understanding of immune dysregulation in PD. This work may ultimately help facilitate the development of targeted immunomodulatory therapeutic strategies and biomarker discovery.

## Results

### Systematic survey of literature on peripheral immunity in Parkinson’s disease

To ensure a comprehensive and unbiased assessment of existing research on peripheral immune cell alterations in PD, a scoping review was conducted in accordance with the Preferred Reporting Items for Systematic reviews and Meta-Analyses extension for Scoping Reviews (PRISMA-ScR) guidelines (Fig. 1A). A predefined search strategy was applied to the PubMed database (Table S1), and all retrieved records underwent title and abstract screening by two reviewers. Eligibility was determined using prespecified inclusion criteria. As such, studies reporting only inflammatory mediators in biofluids (*e.g.,* plasma/serum, CSF, saliva, tear or exosomes), studies pertaining exclusively on the microbiome, and studies of other synucleinopathies (*e.g.,* dementia with Lewy Bodies, multiple system atrophy) were excluded. At level-1 screening, we also excluded papers describing non-human primary cells or CNS-resident immune cells and glia. Moreover, full-texts were assessed, and we excluded any reviews, meta-analyses, clinical trials, non-English articles, and records without easily accessible full-texts. Following expert review by five principal investigators, we further excluded studies lacking matched healthy controls, as well as genome-wide association studies (GWAS), and studies focused solely on platelets or red blood cells. Nineteen additional studies were identified through reference screening and expert recommendation. Ultimately, a total of 285 articles were included into the qualitative synthesis, encompassing patient demographics, methodological characteristics, study design features, and a summary of principal findings.

**Figure 1:**
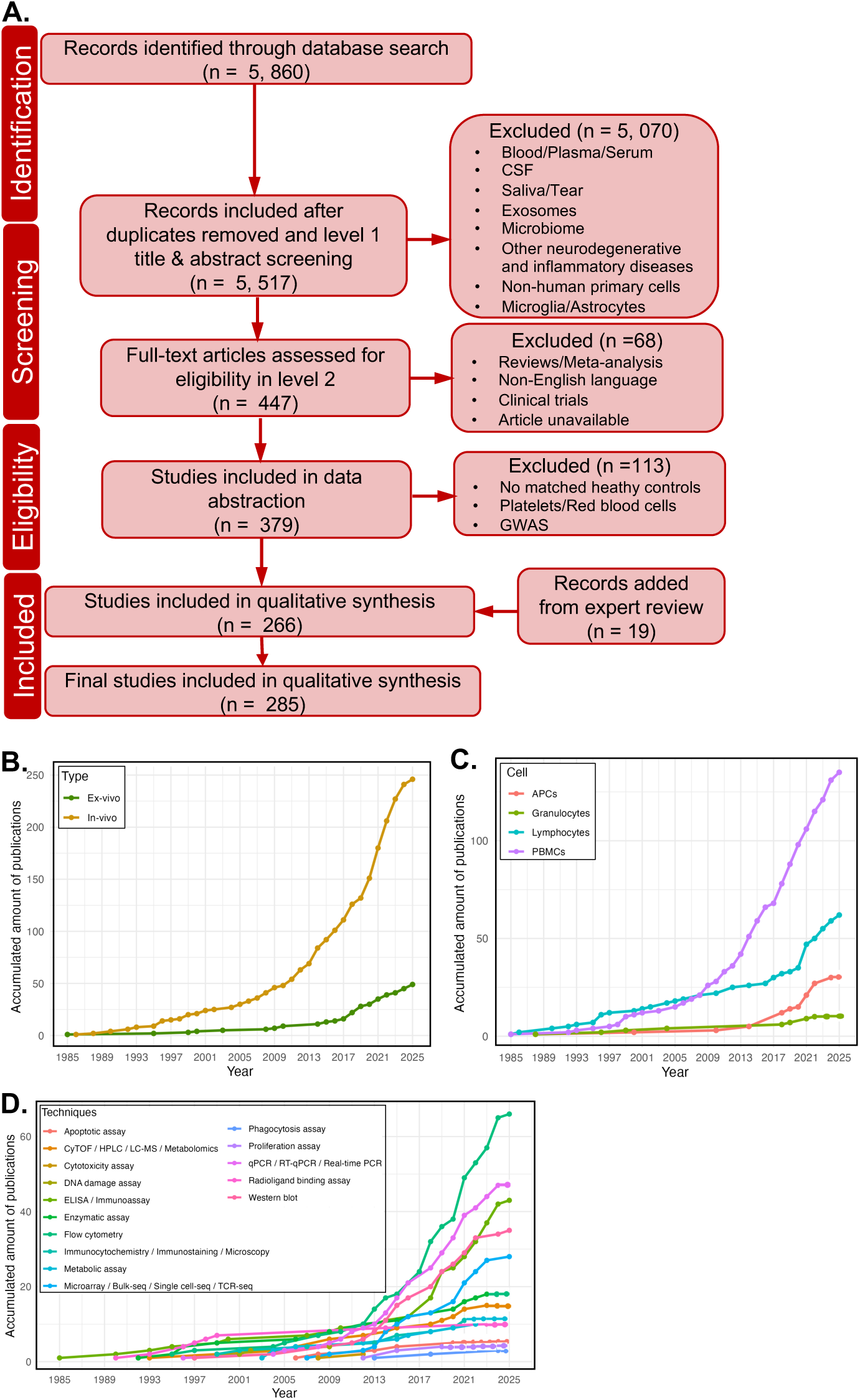
Longitudinal overview of the growth, scope, and methodological evolution of peripheral immune cell phenotyping in Parkinson’s disease. **A.** Preferred Reporting Items for Systematic reviews and Meta-Analyses extension for Scoping Reviews (PRISMA-ScR) flow diagram summarizing the identification, screening, eligibility assessment, and inclusion of peripheral immune cell studies in Parkinson’s disease (PD). **B.** Cumulative number of publications over time stratified by study type, distinguishing *in vivo* analyses of immune cells directly isolated from blood and *ex vivo* studies assessing stimulus-evoked activation and resolution responses of PD immune cells in culture. **C.** Temporal trends in cumulative publications highlighting a progressive shift toward cell-type-resolved analyses of PD peripheral immune populations. **D.** Cumulative adoption of experimental and analytical techniques used for immune cell profiling in PD, illustrating the transition from early single-analyte immunoassays and targeted biochemical approaches to diversified functional assays and high-dimensional methods. PBMCs, peripheral blood mononuclear cells; APCs, antigen-presenting cells.

Peripheral immune cell profiling in PD first gained momentum in the late 20^th^ century (Fig. 1B, C), primarily through single-analyte antibody-based immunoassays (*e.g.,* ELISA, ELISpot) and targeted biochemical techniques to evaluate gene/protein expression, enzymatic activity, metabolism, and DNA damage. (Fig. 1D). These early studies focused on broad phenotyping of peripheral blood mononuclear cells (PBMCs) which include lymphocytes (T-cells, B-cells and natural killer (NK) cells) (Fig. 1C). The field expanded steadily through the 2000s, with approximately 50 primary studies published by 2010 that described alterations in the proportions and “*states*” of circulating immune cell subsets isolated directly from blood of PwPD (referred to as *in vivo* studies) (Fig. 1B). From 2010 onwards, there was a marked acceleration in the number and scope of studies, reflecting both methodological advances and growing recognition of immune contributions to PD. Research increasingly incorporated cell type-specific analyses of positively or negatively selected subtypes of blood-derived lymphocytes or myeloid cells, including antigen-presenting cells (APCs) and granulocytes as opposed to bulk PBMCs (Fig. 1C). During this period, a greater number of functional studies also emerged, employing *ex vivo* cell culture systems to assess immune “*traits*” defined by stimulus-evoked responses (referred to as *ex vivo* studies) (Fig. 1B). In parallel, clinical studies using easily accessible methods to quantify major immune populations in the blood (referred to as *rudimentary immunophenotyping*), such as basic flow cytometry (based on cell shape and size), became more common, facilitating translational applications. Rudimentary immunophenotyping has identified reproducible changes in immune cell populations and markers for inflammation in peripheral blood from PwPD (see Table 1 for the list of included studies). Most notable are increases in the neutrophil to lymphocyte ratio (NLR), with ten papers (out of 14 studies that measured NLR) reporting this change relative to healthy donors (Fig. S1).

**Table 1:**
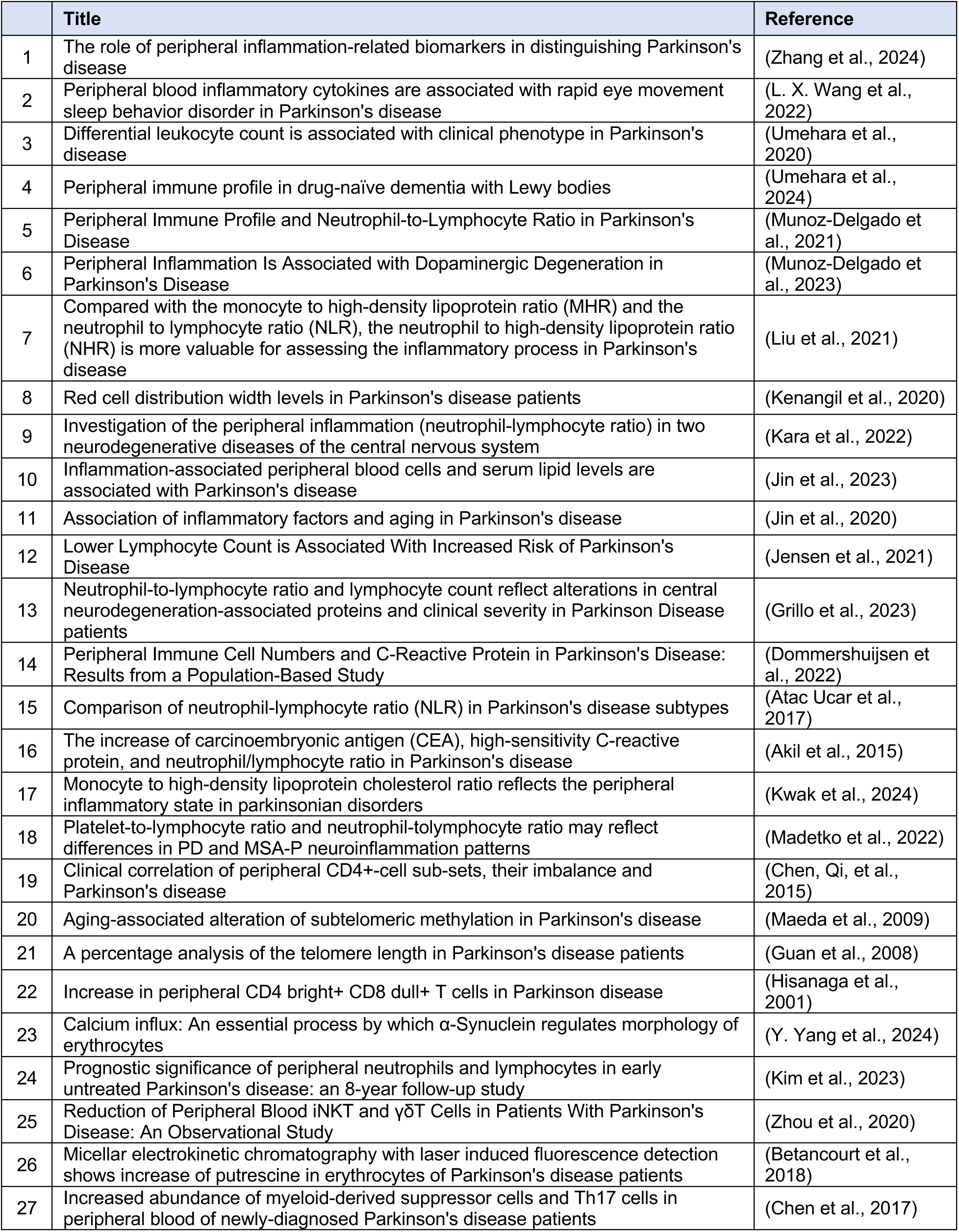
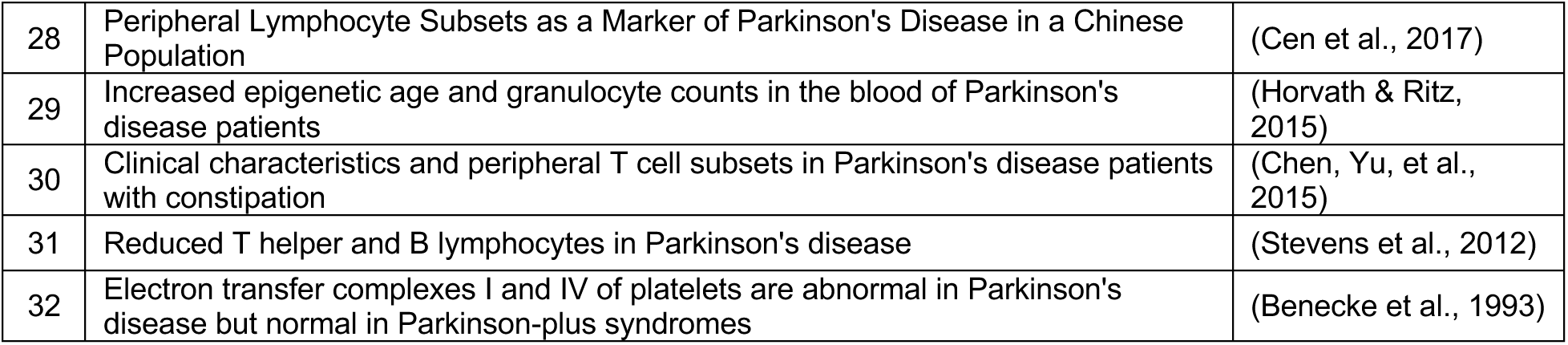
List of studies included as *Rudimentary immunophenotyping*.

Technological diversification was a defining feature of the past 15 years (Fig. 1D). Classical immunoassays were complemented by approaches interrogating cellular functions, such as phagocytosis, proliferation, apoptosis and cytotoxicity, as well as unbiased large-scale methodologies, including bulk and single-cell sequencing. Despite this expansion, flow cytometry has remained the primary tool for immune cell profiling in PD, highlighting its central role in the field. Taken together, this longitudinal overview reveals a steady and then rapid growth of research examining peripheral immune cell alterations in PD. The evolution of methodologies has enabled increasingly detailed characterization of immune phenotypes and functions. Synthesizing these developments is essential to identify key findings, methodological disparities, and remaining knowledge gaps.

### Synthesis of demographic and clinical heterogeneity across immune-focused PD studies

Differences in cohort composition, including sample size, sex distribution, age, disease stage/severity, clinical assessments, therapies, and exclusion criteria can influence immune measurements and may partly explain the disparate findings reported in the PD literature. By examining these parameters, we aimed to provide a clearer contextual framework for interpreting immune alterations across studies stratified by study types (*in vivo* versus *ex vivo*) (Fig. 2).

**Figure 2:**
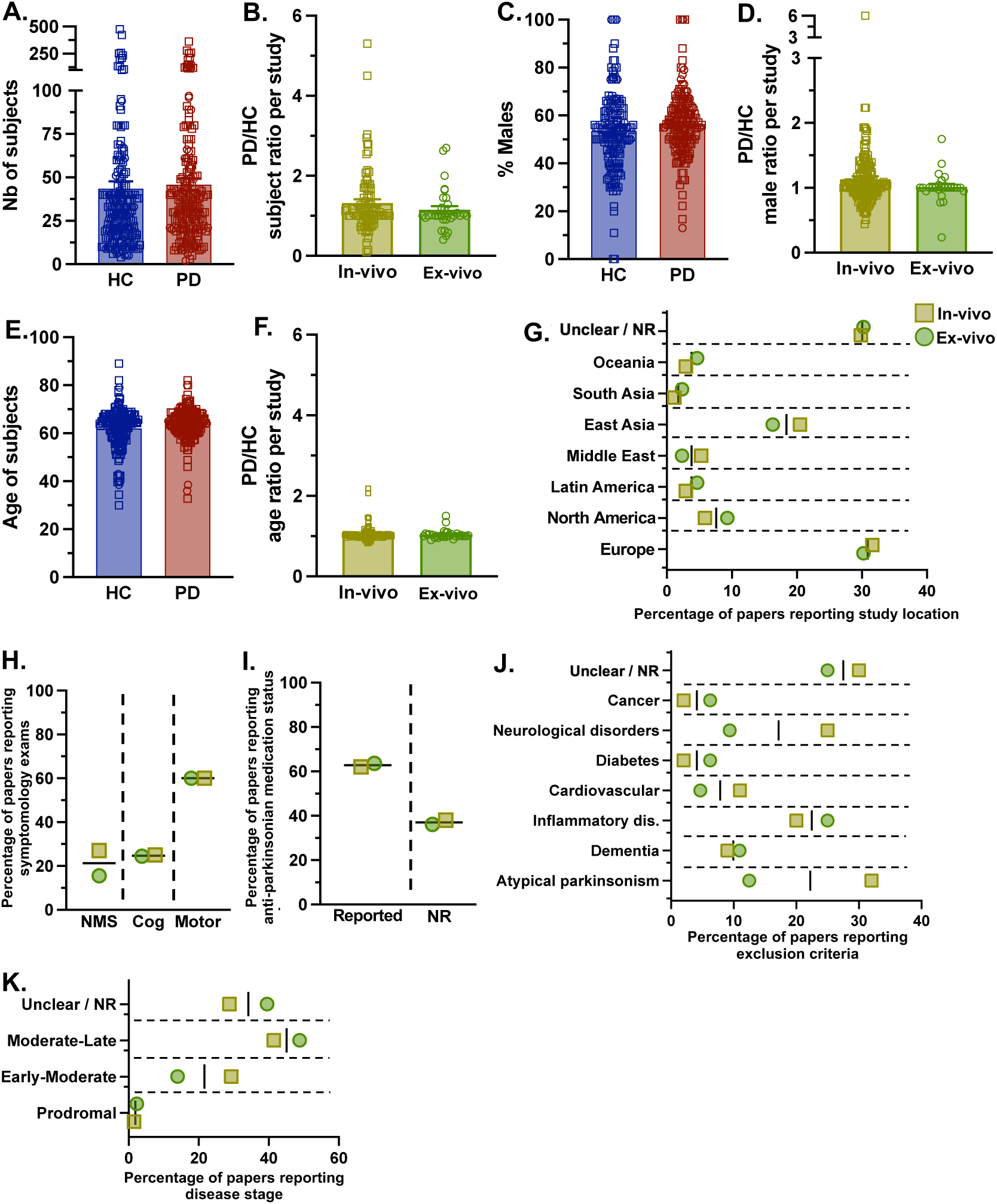
Participant demographics and clinical variables in peripheral immune studies of Parkinson’s disease. A-F. Distributions and ratios of cohort size, sex, and age between healthy control (HC) and PD groups across included *in vivo* and *ex vivo* studies. **G-K.** Frequency of publications reporting geographic and ethnic origins of study cohorts, clinical symptom assessments, anti-parkinsonian medication status, exclusion criteria and disease stage. NR, not reported; NMS, non-motor symptoms; Cog, cognitive.

Analysis of 285 studies (see Tables 2-4 for the list of included studies) revealed substantial variability in both cohort characteristics and the extent of reporting. Overall, study sizes spanned a broad range (^5 to]400 subjects), but the numbers of participants in PD and healthy control (HC) groups were generally comparable across study types (Fig. 2A, B). Sex distribution and age were similarly matched between cohorts, with an average male proportion of 53% in HC and 56% in PD groups (Fig. 2C, D), and mean ages of 62 and 64 years, respectively (Fig. 2E, F). However, the sex balance between the groups within studies was not always achieved with nearly 46% of *in vivo* studies having more males in the PD group (Fig. 2D). These studies have also enrolled participants from varying ethnic origins (Fig. 2G), with ∼30% being European descents, ∼20% from East Asian countries, and 10% from North America. While less than 5% of recruited subjects are either from Latin America, Middle East, South Asia or Oceania. Of note, about a third of studies did not report the ethnicities of their participants.

**Table 2A:**
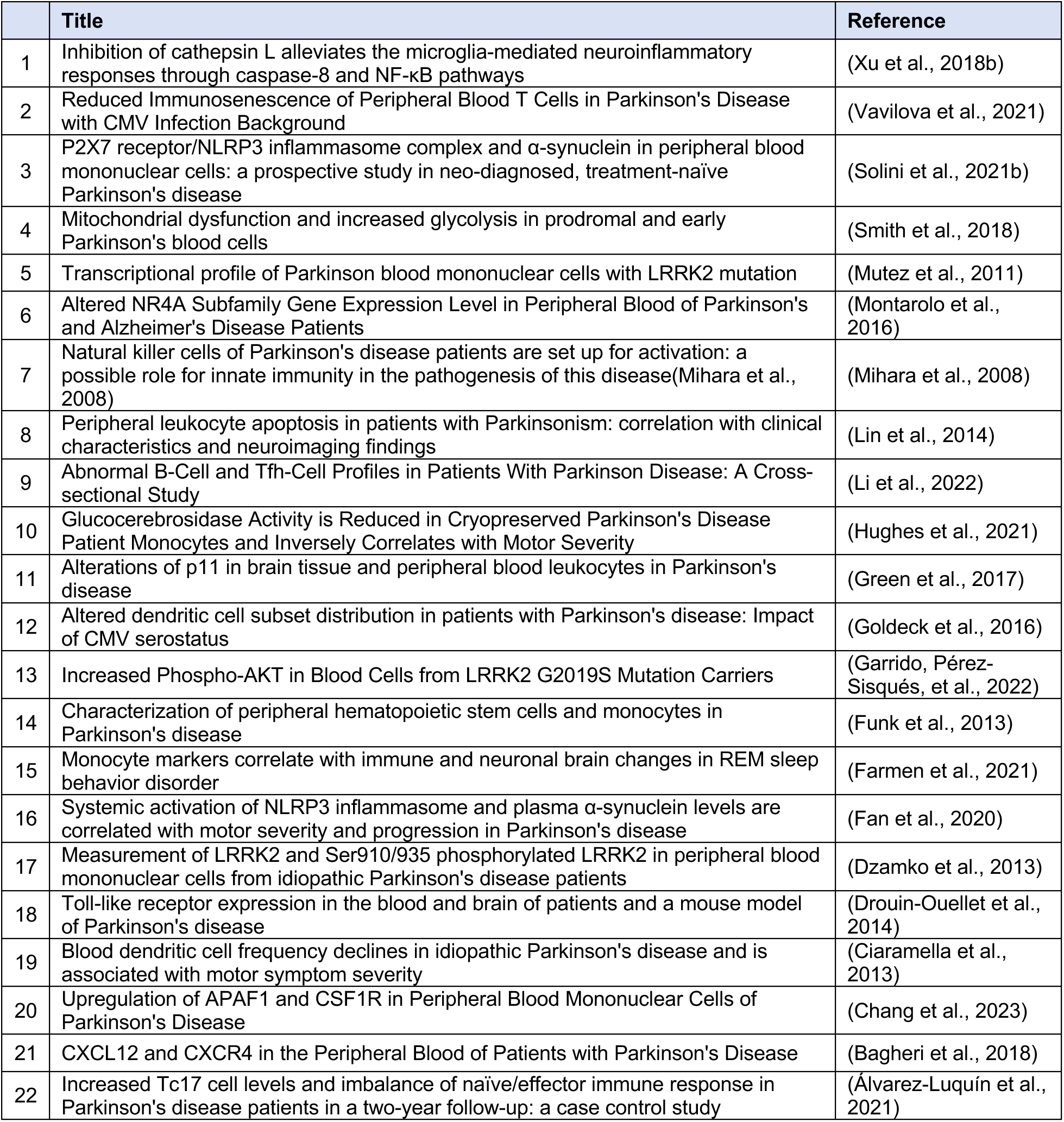
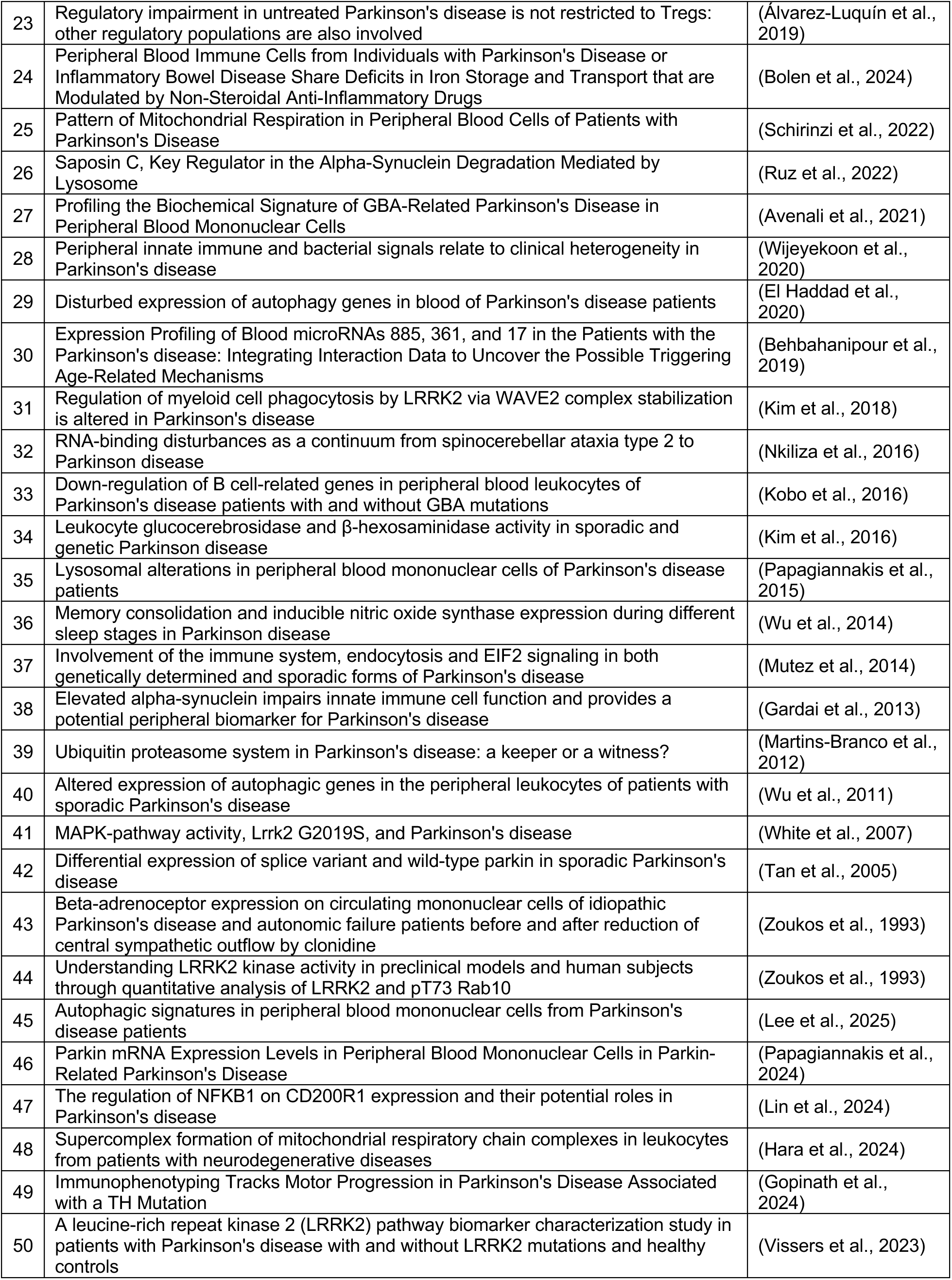
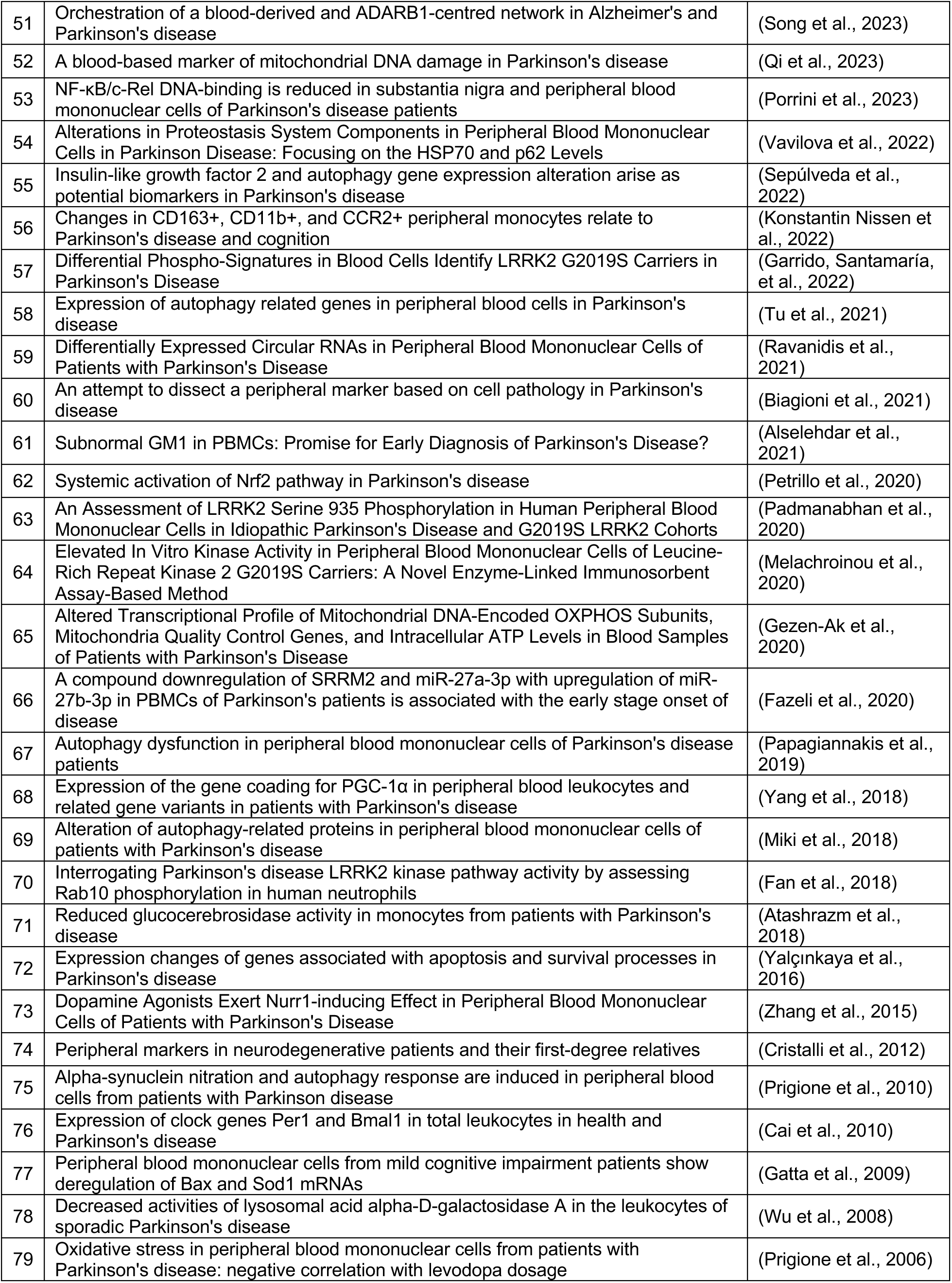
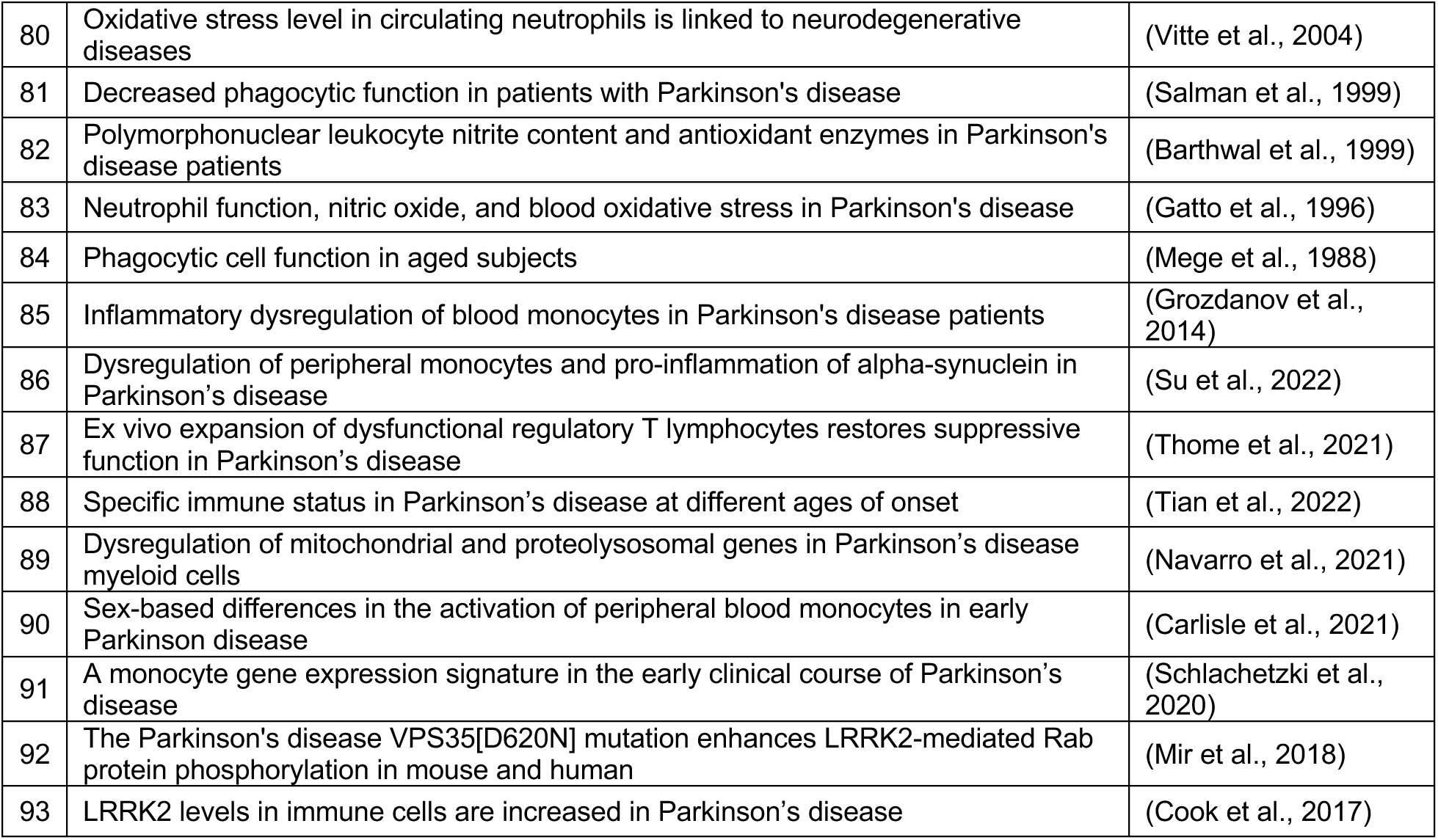
List of studies included as *In vivo advanced immunophenotyping* (innate)

**Table 2B:**
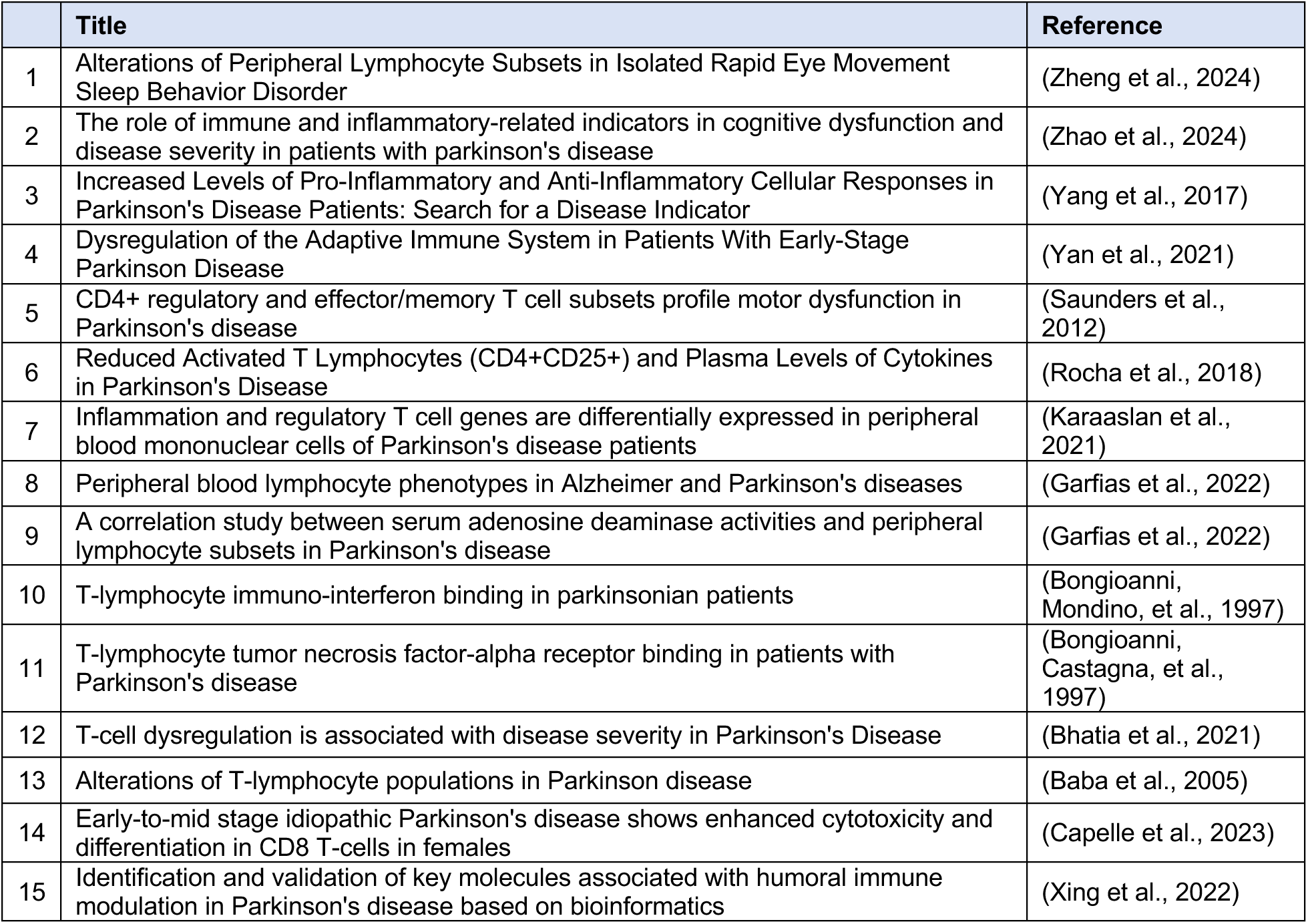
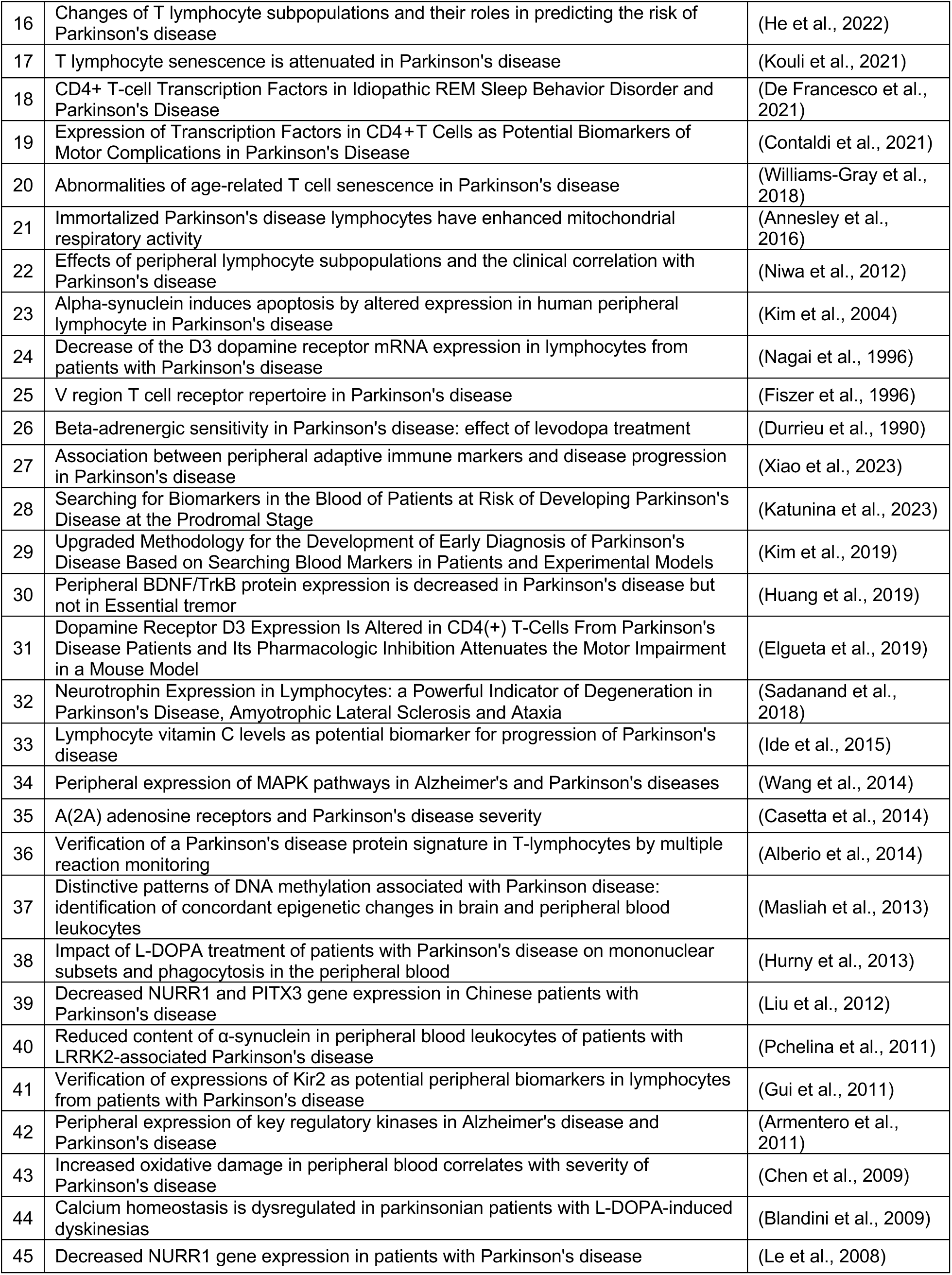
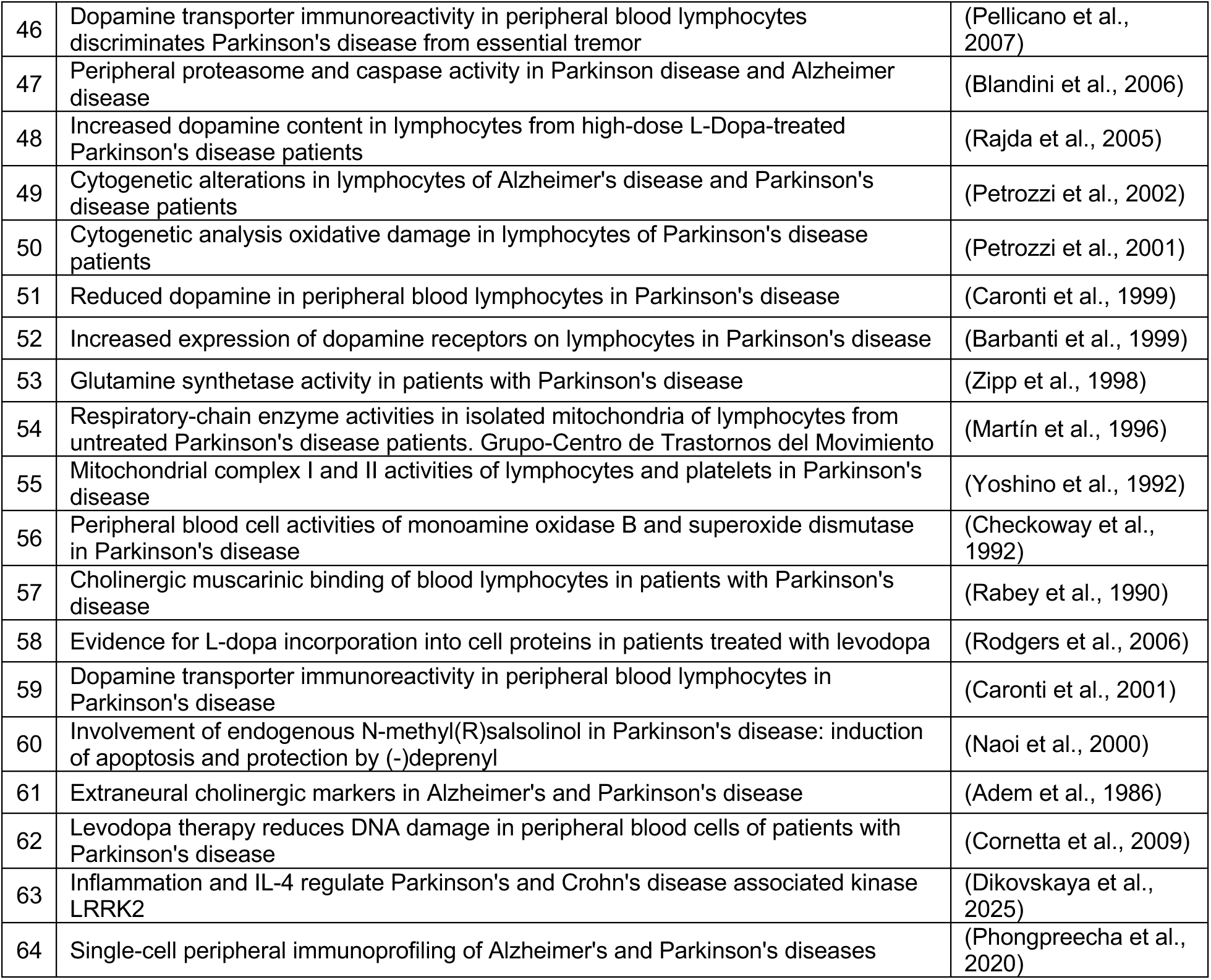
List of studies included as *In vivo advanced immunophenotyping* (adaptive)

**Table 3:**
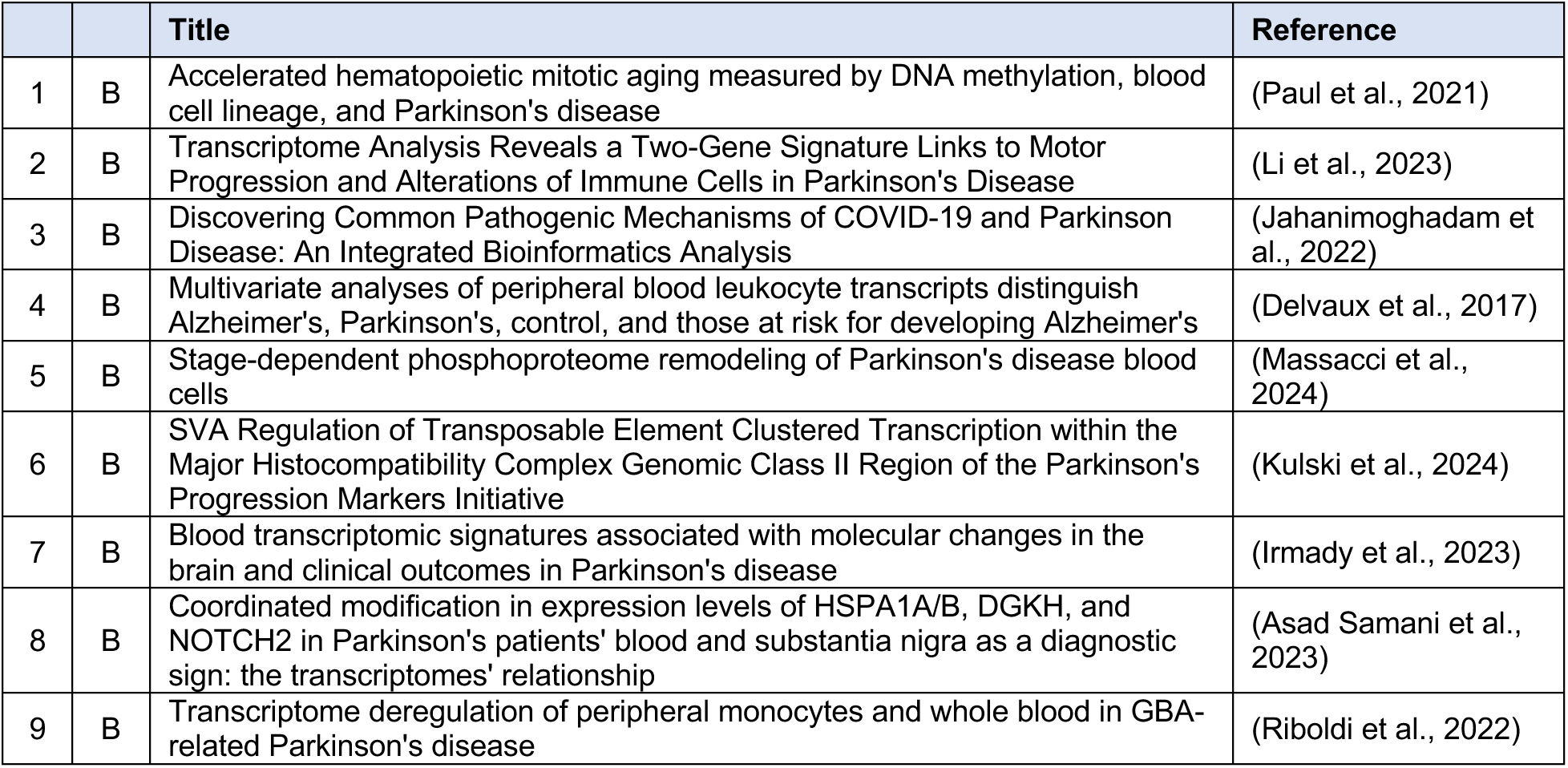
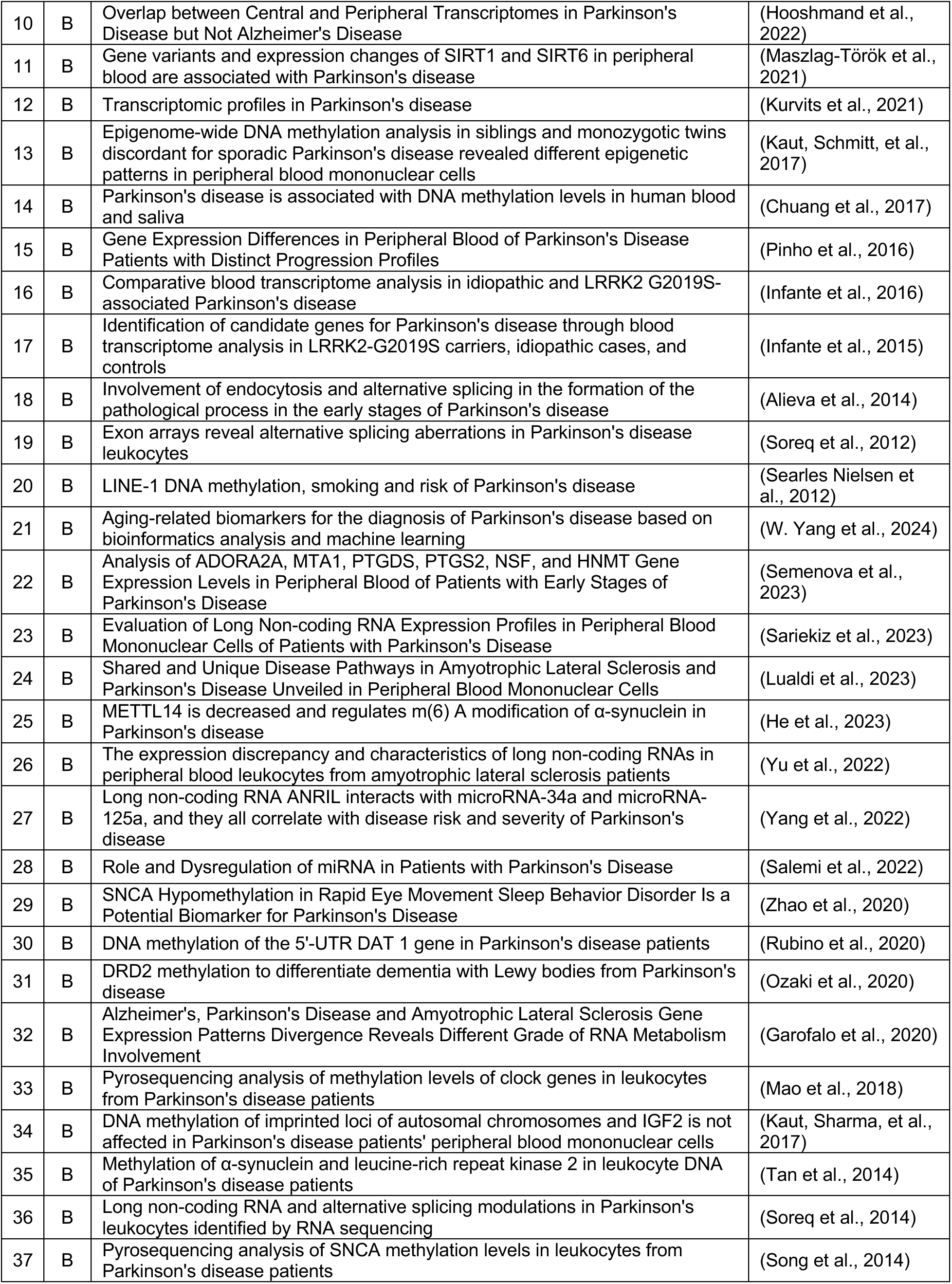
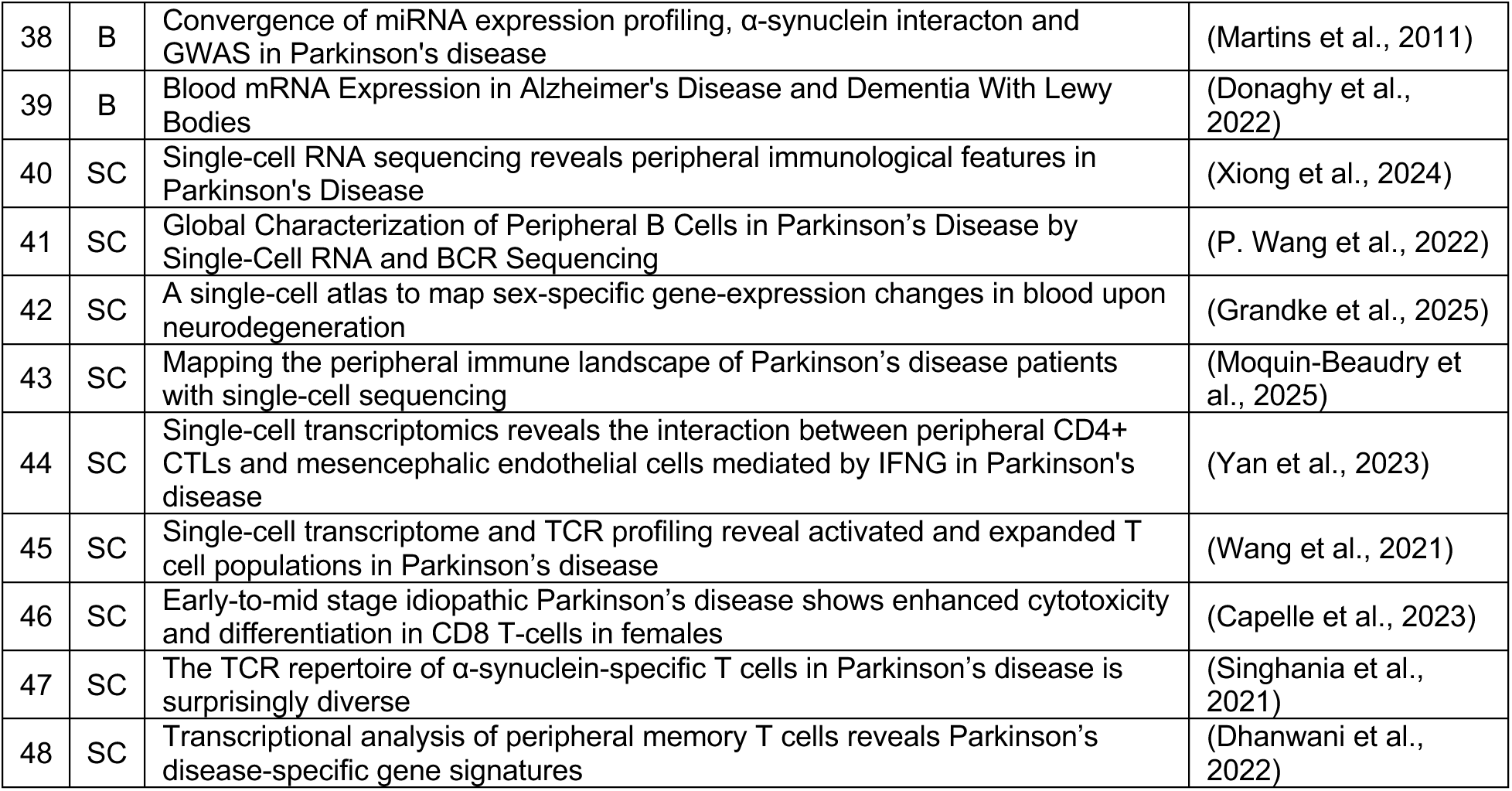
List of studies included as *Sequencing-based immunophenotyping*.

**Table 4:**
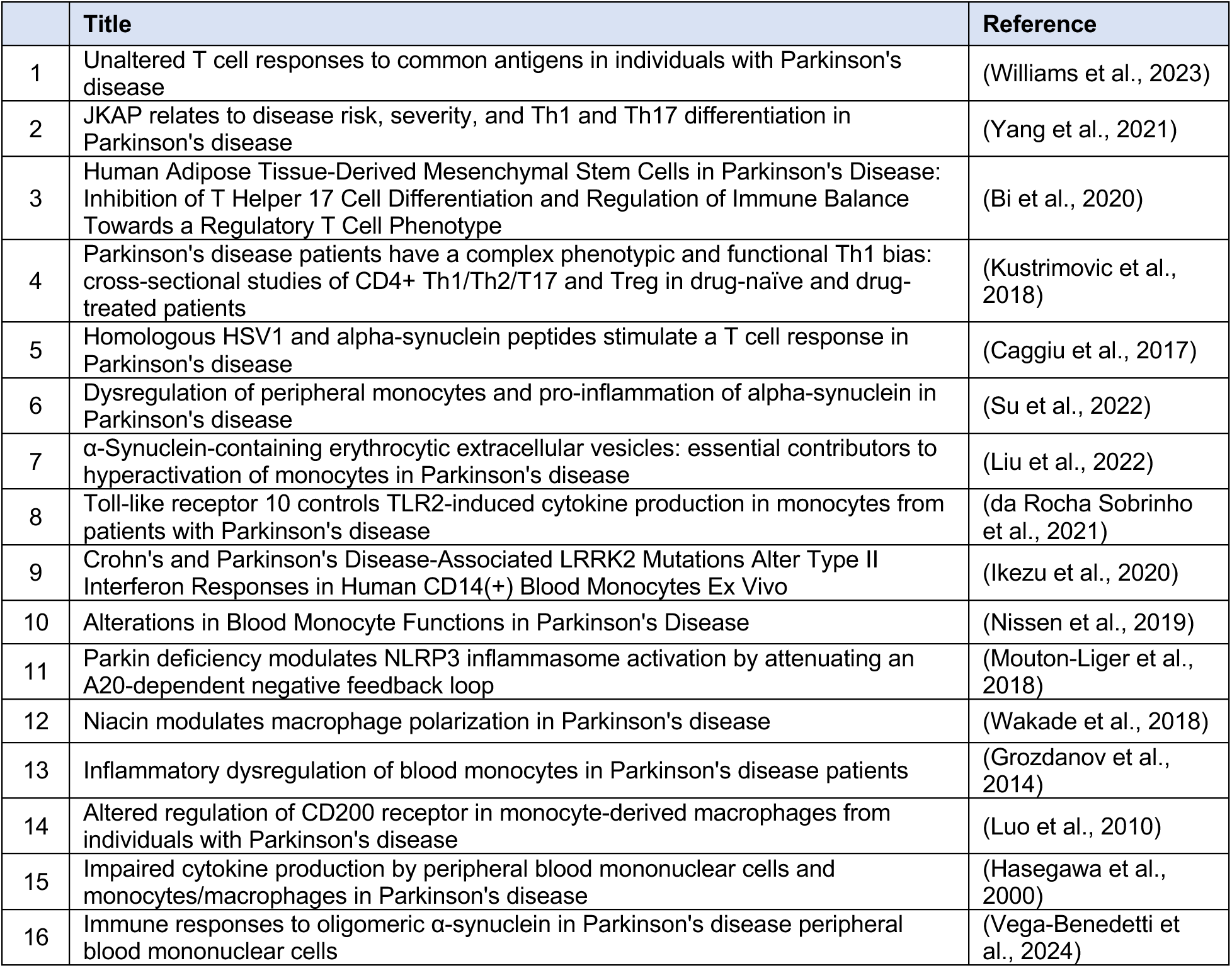
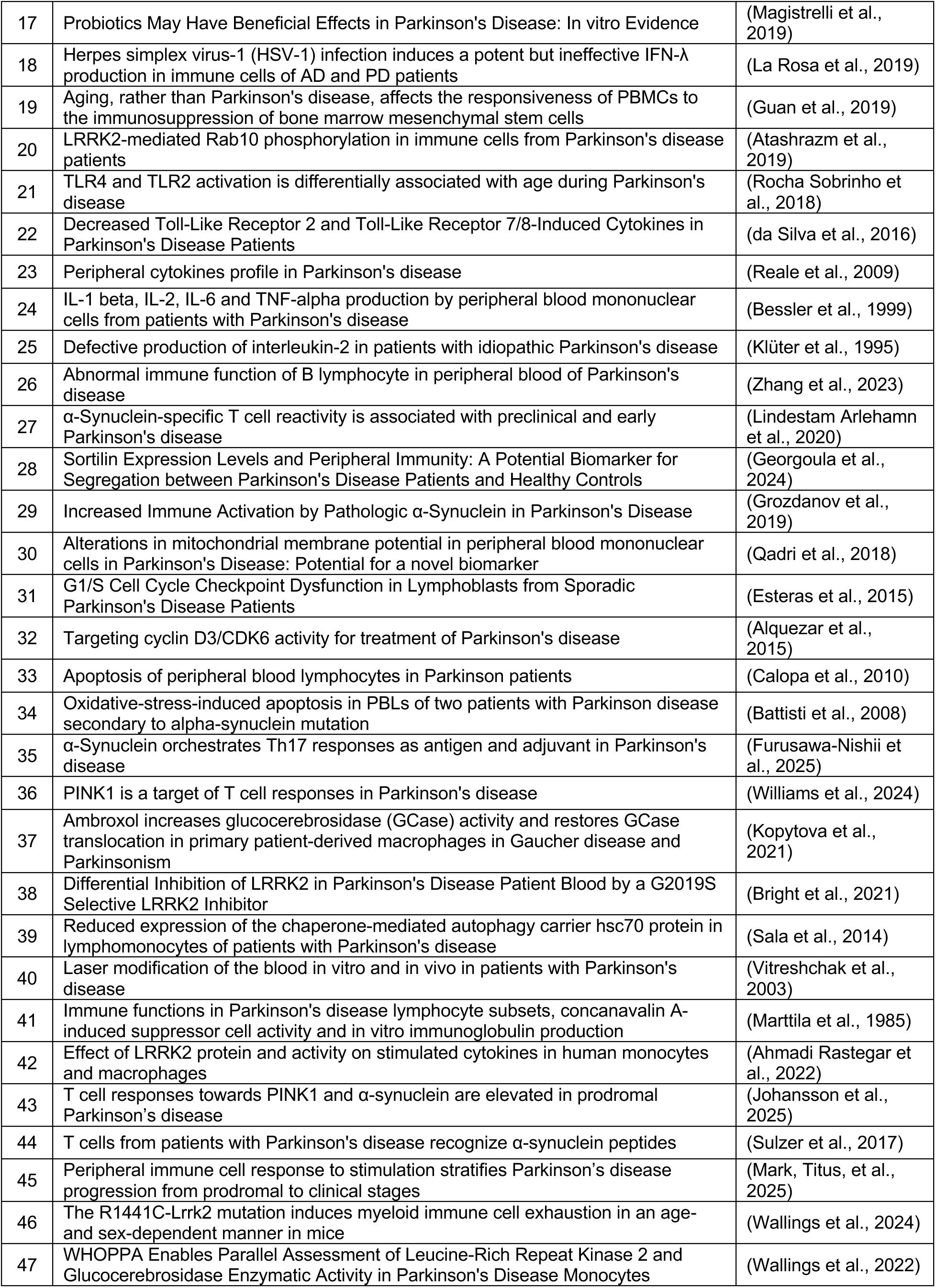
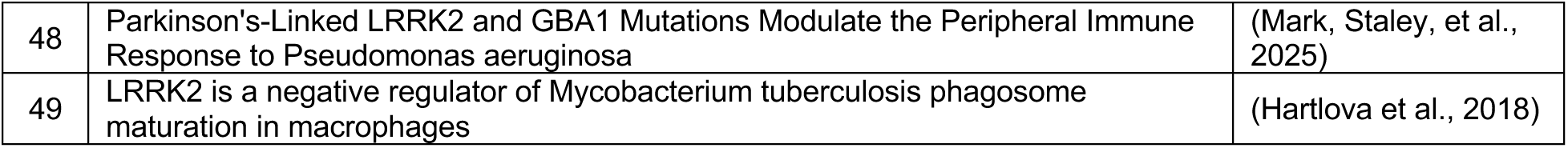
List of studies included as *Ex vivo immunophenotyping*.

Additionally, clinical assessments were inconsistently documented. Although ∼60% of studies reported motor symptom evaluations, much fewer included details of non-motor or cognitive assessments (∼20%) (Fig. 2H). Reporting of anti-parkinsonian medication status was also incomplete, with nearly 40% of studies failing to specify treatment details (Fig. 2I). Furthermore, exclusion criteria exhibited marked variability (Fig. 2J); while atypical parkinsonism, dementia, inflammatory disorders, cardiovascular disease, diabetes, neurological comorbidities, and cancer were noted to different degrees, about a quarter of studies listed no exclusion criteria. These results were largely comparable across *in vivo* and *ex vivo* studies, with a notable exception that neurological disorders and atypical parkinsonisms were more frequently excluded in *in vivo* studies. Beyond these factors, disease stage/severity was also unevenly captured across studies. Cohorts ranged from prodromal individuals (diagnosed with REM sleep behaviour disorder, RBD) (<2%) to those with early-to-moderate PD (∼20%) and moderate-to-late PD (∼45%), whereas a sizeable proportion of studies provided unclear or no information on disease stage/severity (∼35%) (Fig. 2K). Of note, for our analysis, and in keeping with the literature, we categorized patients with “early to moderate stages” as patients with disease duration <5 years and/or manifesting symptoms/signs observed early in the disease according to established clinical exams, such as H&Y scales I/II, Webster ratings <8, Braak stages I/II or UPDRS-III ratings <18. Within each category, the percentage of participants recruited varied substantially, reflecting inconsistent sampling strategies and highlighting another dimension of clinical heterogeneity that may influence immune readouts. Overall, these data suggest that the field has achieved consistency in reporting and matching age and sex across cohorts, although remains variable in documenting ethnicity, disease stage/severity, treatment status, and exclusion criteria. This uneven reporting reveals persistent gaps in standardization in peripheral immune research in PD.

### Overview of in vivo studies illustrating peripheral immune states in PD

To characterize methodological trends in peripheral immune cell profiling in *in vivo* studies, we examined key steps from sample acquisition to data analysis across 157 PD studies. These were stratified according to the primary immune populations analyzed – innate or adaptive (see Tables 2A and 2B for the list of included studies). Notably, although 100 studies assessed both innate and adaptive immune cells, they were categorized based on the immune cell type(s) emphasized in the major conclusions. PBMCs were typically obtained by density-gradient centrifugation, aliquoted, and either analysed immediately (fresh) or cryopreserved (frozen) for future downstream assays (Fig. 3A). Although anticoagulants are required to preserve cell integrity and ensure successful isolation of PBMCs, their use can influence immune function. We thus compared the choice of anticoagulants reported in *in vivo* studies. Most reports did not specify the reagent used regardless of immune cell types assessed, while EDTA was the most common among those that did (Fig. 3B). The use of fresh *versus* cryopreserved PBMCs was also evaluated. *Adaptive* studies more often relied on freshly processed samples, while *innate* studies showed roughly equal distribution between frozen and fresh material, with ∼10% of all studies lacking sufficient information to determine sample state (Fig. 3C). Reporting of centrifugation conditions across both density-gradient separation and subsequent wash steps showed that most papers documented g-force rather than revolutions per minute (RPM) and over half did not fully report spin parameters (Fig. 3D). Moreover, despite the potential impact of storage conditions and preparation steps on cell viability and yield, more than half of the studies omitted or incompletely described these features (Fig. 3E), which impairs the evaluation of the cell quality, and thus the data obtained, during the analysis. The importance of ensuring the processing pipeline is operator and site independent cannot be underscored enough. Several groups have indeed investigated the extent to which this could be achieved by deep immunophenotyping control PBMCs at two different sites (Wallings et al., 2022) followed up by *ex vivo* studies on immune inducibility of idiopathic and genetic PD (LRRK2 and GBA1 variants) PBMCs from PD cryorepositories in response to inflammatory stimuli or pathogens (Hughes et al., 2025; Mark, Staley, et al., 2025).

**Figure 3:**
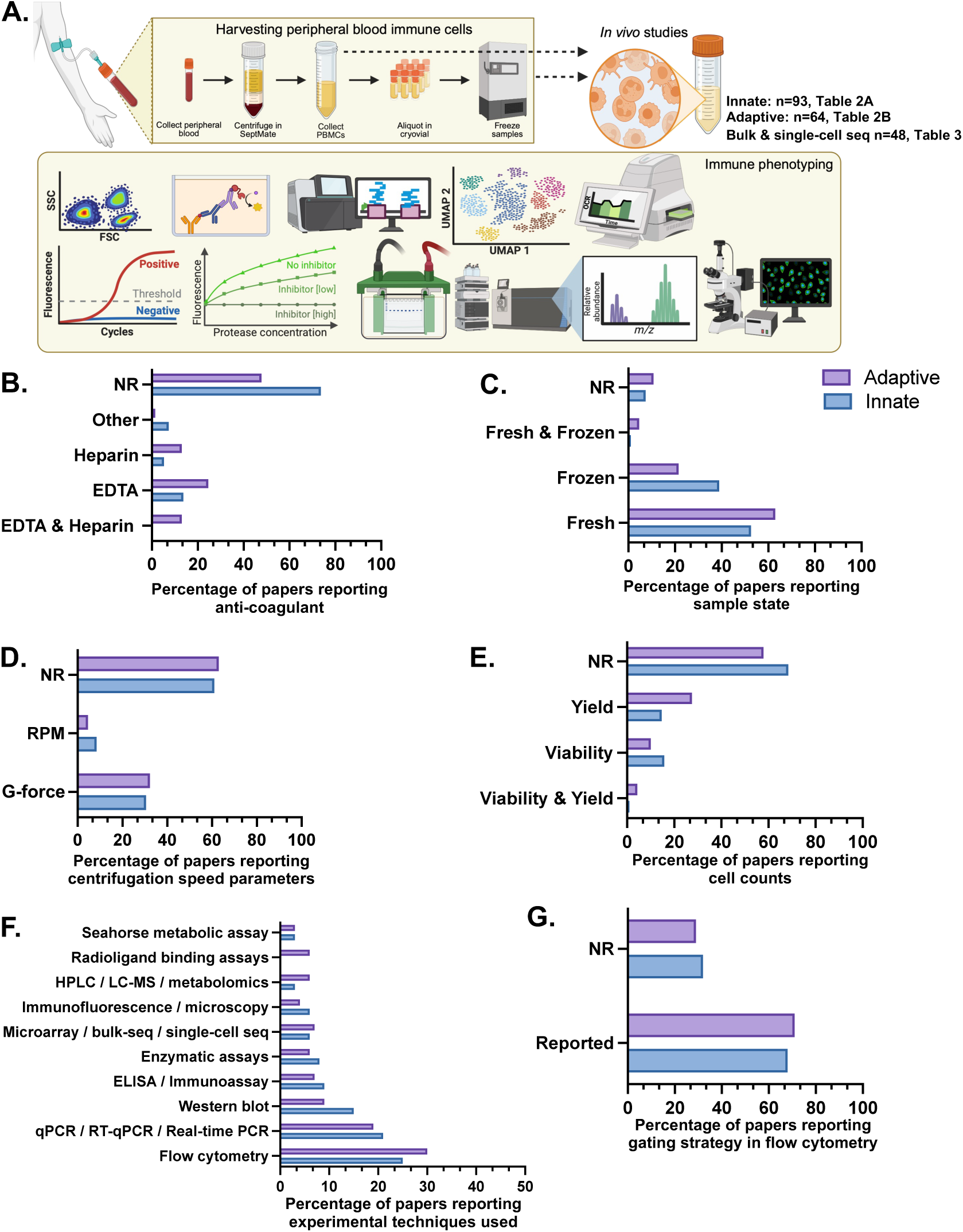
Methodological landscape of *in vivo* peripheral immune cell profiling in Parkinson’s disease. **A.** Schematic representation of experimental workflows in *in vivo* PD peripheral immune research, spanning blood collection, PBMCs isolation, and downstream immune phenotyping using molecular, biochemical, cytometric, metabolomic, imaging-based and sequencing approaches. The number and associated list of studies interrogating innate, adaptive, or sequencing-based immune phenotypes are indicated. **B-G.** Reporting of anticoagulants for blood collection, sample handling practices, centrifugation parameters for PBMC isolation, cell yield and viability measurements, experimental platforms for immune characterization, and gating strategies in flow cytometry-based *in vivo* PD studies, stratified by immune population focus (innate *versus* adaptive). RPM, revolutions per minute (RPM); G, g-force.

We also noted a wide array of analytical platforms that was applied to assess immune *states*. Flow cytometry was the dominant approach, followed by qPCR, Western blotting, and ELISA performed on immune cell lysates and/or supernatants, whereas advanced proteomic, metabolic or imaging assays were less commonly utilized (Fig. 3F). Among flow cytometric studies, documentation of gating strategies was inconsistent, with roughly one-third of *in vivo* studies failing to present gating profiles (Fig. 3G). Furthermore, studies varied in gating strategies. For instance, in the case of innate cells, some use live/dead discrimination followed by size and granularity parameters for the inclusion gate, whereas others define gates using myeloid-specific cell surface markers such as CD64, TLR2, or CD45 combined with negative selection for markers like CD3, CD19, CD56, and CD66b. Importantly, certain molecules used for classification are also subject to proteolytic cleavage from the surface, as such disparities in protocol or antibodies might result in differences in detected proteins and definition of subpopulations.

In addition to hypothesis-driven methods, 48 studies implemented unbiased, sequencing-based immune profiling, including bulk RNA-sequencing (RNAseq), long non-coding RNA sequencing (lncRNA-seq), epigenomic and proteomic analyses, as well as single-cell RNAseq, with four of these studies also incorporated T-cell and B-cell receptor sequencing (TCR/BCR-seq) (Fig. 3F, 4A top panel; see Table 3 for the list of included studies). Among the nine single-cell RNA-seq studies, we extracted the number of cells analyzed and the quality-control parameters applied (Fig. 4A, bottom panel) as the rigor and interpretability of sequencing studies depend in part on sampling depth and preprocessing strategies. We observed considerable variability in cell counts, with most studies analyzing between 10,000 and 100,000 cells, and only two exceeding 100,000 cells. In contrast to this heterogeneity in sampling depth, preprocessing practices were comparatively consistent, with seven of nine studies reporting the application of standard quality-control measures including removal of low-quality cells based on mitochondrial transcript reads (typically > 5-20%) and doublet detection using tools such as scDblFinder or DoubletFinder. In droplet-based single-cell RNAseq experiments, background contamination from ambient RNA is common, arising from both encapsulated cells and extracellular RNA released during cell lysis. Nevertheless, only two studies reported correcting for ambient RNA contamination using SoupX. To summarize cell type-specific aberrant pathways, we extracted major GO and KEGG biological processes reported in nine single-cell RNAseq studies. Altogether, 118 pathways were described as upregulated and 9 as downregulated in PD compared to matched controls. We then compiled dysregulated pathways denoted in at least two independent studies and mapped to specific immune cell types, including classical monocytes, cytotoxic NK cells, mucosal-associated invariant T-cells (MAIT), CD4+ and CD8+ effector T cells (CTLs), CD4+ and CD8+ memory T cells (TEM/TEMRA), and B-cells (Fig. 4B). Among the shared altered biological processes, innate immune response, T-cell receptor signaling/activation, and leukocyte migration/chemotaxis were repeatedly reported within the same cell types (Capelle et al., 2023; Dhanwani et al., 2022; Moquin-Beaudry et al., 2025; P. Wang et al., 2022; Wang et al., 2021; Xiong et al., 2024; Yan et al., 2023). Notably, cytotoxic NK and CD8+ TEM/TEMRA populations were most frequently implicated, with convergent evidence pointing toward heightened activation states in PD. Next, we synthesized immune *state*-level reporting across *in vivo* studies to evaluate how often PD-associated changes were described in terms of cell proportions and/or molecular features, encompassing inflammatory gene/protein expression, cell surface receptors/transporters, oxidative/mitochondrial stress, phagocytosis/autophagy-lysosomal axis and cell death, and stratified by study types (Fig. 4C). Within the innate immune compartment (see Table 2A for the list of included studies), the monocyte frequencies were often unchanged (12 out of 24) although few studies reported shifts within monocyte subsets, most commonly both an increase and a reduction of classical (CD14^high^CD16^-^) and intermediate subsets (CD14^low^CD16^-^) accompanied by an expansion of non-classical monocytes (CD14^-^CD16^+^) (Fig. 4D). Beyond monocyte population, the changes in the cellular composition of other innate cells are less studied, with one study reporting a reduction in frequencies of both myeloid and plasmacytoid dendritic cells (m-and p-DCs), particularly in newly diagnosed patients (Ciaramella et al., 2013). While another showed that cytomegalovirus status in PD affects mDC subpopulations (Goldeck et al., 2016), highlighting infections as environmental modulators of PD-immunity. Moreover, the finding that higher proportions of cytotoxic NK cells are described in PD *versus* HC is reproduced in five *in vivo* studies (Fig. 4D, F).

**Figure 4:**
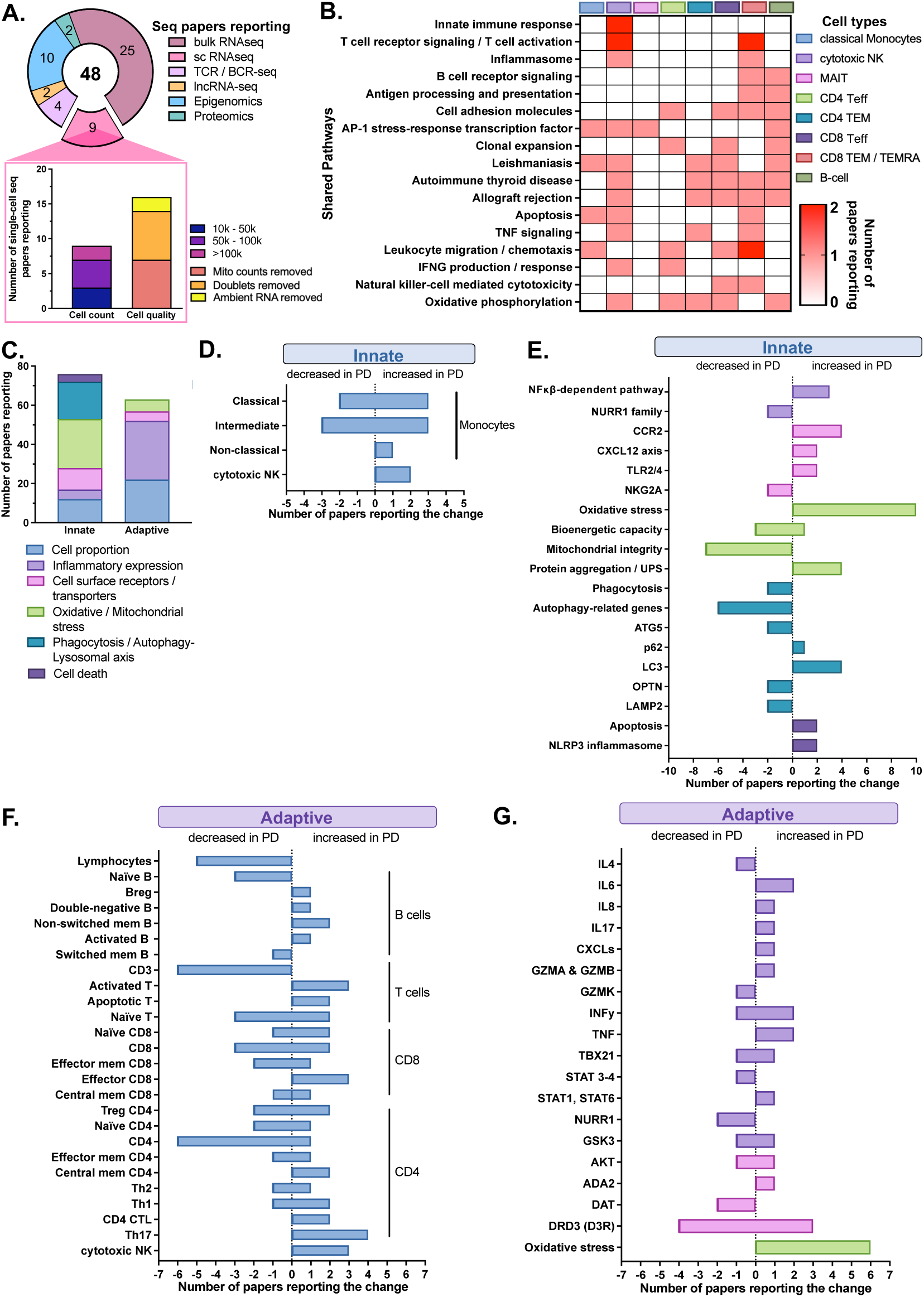
Peripheral immune states uncovered by *in vivo* phenotyping in Parkinson’s disease. **A.** Pie chart depicting the total number of studies employing sequencing-based immune profiling of PBMCs in PD, stratified by omics modality. Shown in the inset bar graph are the methodological features of the nine single-cell RNA sequencing (RNAseq) studies, including the number of cells analyzed (stratified as 10,000-50,000; 50,000-100,000; and >100,000 cells) and reported quality-control measures (removal of cells with high mitochondrial transcript content, doublet detection, and ambient RNA correction). **B.** Heatmap summarizing shared GO/KEGG pathways described as dysregulated in specific immune cell populations across published single-cell RNAseq studies. Only pathways noted in at least two independent studies are presented; color intensity reflects the number of studies reporting each pathway within a given cell type. **C.** Synthesis of the number of *in vivo* publications describing PD-associated alterations in terms of cell proportions and distinct molecular features, stratified by innate and adaptive immune populations. **D-G.** Reporting of directionality and frequency of reported changes in cell proportions and molecular features of either innate or adaptive immune cell subsets in PD. RNA sequencing, RNAseq; TCR, T-cell receptor; BCR, B-cell receptor; lncRNA, long non-coding RNA; NK, natural killer; MAIT, mucosal-associated invariant T cells; Teff, effector T cells; TEM, effector memory T cells; TEMRA, terminally differentiated effector memory cells re-expressing CD45RA; IFNG, interferon gamma; TNF, tumor necrosis factor; CCR2, chemokine receptor 2; iNOS, inducible nitric oxide synthase; COX2, cyclooxygenase-2; NF-κB, nuclear factor kappa-light-chain-enhancer of activated B cells; NURR1, nuclear receptor-related 1; CXCLs, CXC motif chemokines; TLR2/4, toll-like-receptors 2 & 4; NKG2A, killer cell lectin-like receptor C1 or KLRC1; UPS, ubiquitin-proteasome system; ATG5, autophagy-related protein 5; p62, sequestosome-1 or SQSTM1; LC3, microtubule-associated protein 1 light chain 3; OPTN, optineurin; LAMP2, lysosomal-associated membrane protein 2; NLRP3, NOD-, LRR-, and pyrin domain-containing protein 3; Mem, memory; Treg; regulatory T-cells; Th, T helper cells; MDSCs, myeloid-derived suppressor cells; IL, interleukin; TBX21, T-box transcription factor; STAT, signal transducer and activator of transcription; GSK3, glycogen synthase kinase-3; AKT, serine/threonine kinase or PKB; ADA2, adenosine deaminase 2; DAT, dopamine transporter; DRD3 / D3R, dopamine D3 receptor.

These proportional changes were often paralleled by molecular alterations in innate cells that point toward enhanced inflammatory potential and organelle dysfunction, including mitochondria and lysosomes (Fig. 4E). Across eight studies, monocytes from PwPD exhibit higher expression of cell surface of immune sensors previously associated with neurodegeneration (TLR2/4, TREM2), and receptors involved in immune recruitment and infiltration (CXCR4, CCR2 and CD11b), as well as in antigen presentation (HLA-DR) (Fig. 4E; in pink). Three studies also documented increased expression of downstream inflammatory mediators or pathway components, including iNOS, COX-2, TNF and NFκB (Fig. 4E; in light purple), reinforcing a shift toward heightened activation states with possibly involvement of NLRP3-mediated inflammasomes. Notably, although PD innate immune cells seemed dysregulated and prone to enhanced inflammatory responses, these features did not consistently translate into consistent increased secretion of pro-inflammatory cytokines (Fig. 5I, J). This might be explained by several mechanisms, such as compromised vesicular compartment, which may be directly related to a disturbance of the autophagy-lysosomal axis, and/or disturbed energy status, comprising nearly 60% of publications reporting dysregulated molecular features (Fig. 4C; in teal and green, respectively). PBMCs from PwPD exhibit altered expression of key autophagy-lysosomal genes (ATG5, p62, LC3, OPTN and LAMP2) (Fig. 4E; in teal), suggesting an impaired autophagic flux that may hinder chaperone-mediated activities and autophagosome-lysosome fusion leading to the accumulation of autophagic vacuoles. Moreover, lysosomal enzymes have also been investigated in several studies with reports showing either an increase (Cathepsin D/L) or decrease (α-D-galactosidase, glucocerebrosidase, β-galactosidase, Prosaposin) in circulating immune cells from PwPD depending on the proteins being examined and specific cell type (Ruz et al., 2022; Wu et al., 2008; Xu et al., 2018a). Intriguingly, the amounts of certain lysosomal enzymes are found associated with levels of either α-synuclein or inflammatory markers in PD (Avenali et al., 2021; Ruz et al., 2022; Xu et al., 2018a). Additionally, twenty-five studies elucidated that PBMCs from PwPD show evidence of mitochondrial impairments, exemplified by elevated ROS production, reduced bioenergetic capacity and disrupted mitochondrial integrity, including mitochondrial content, SOD activity and mitochondrial supercomplex formation (Fig. 4E; in green). Conversely, the evidence for increased cell death is rather sparse, with four studies involving both apoptotic and NLRP3-mediated pyroptotic pathways (Fig. 4E; in dark purple). Although the impact on cellular viability is less understood, it is important to note that these metabolic alterations could lead to largely different functionalities and efficiencies of innate immune cells.

**Figure 5:**
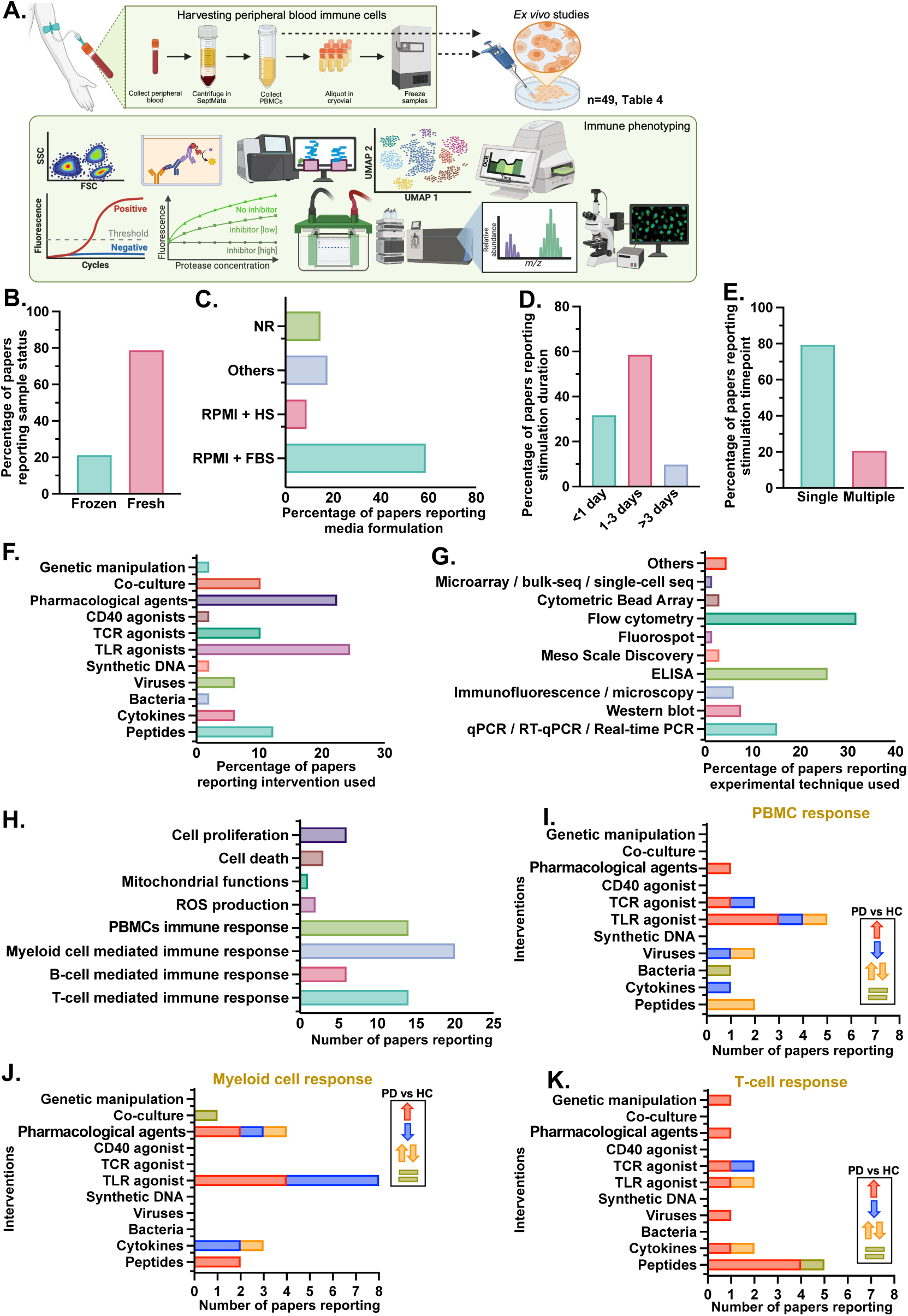
Methodological landscape and immune traits uncovered by *ex vivo* peripheral immune cell profiling in Parkinson’s disease. **A.** Schematic representation of experimental workflows in *ex vivo* PD peripheral immune research, spanning blood collection, PBMCs isolation, cell culture conditions, and downstream immune phenotyping using molecular, biochemical, cytometric, metabolomic and imaging-based approaches. The number and associated list of studies are indicated. **B-G.** Reporting of sample handling practices, culture media formulations, duration and frequency of stimulation, types of intervention and experimental platforms for immune characterization. **H.** Illustration of the number of publications documenting the major functional immune changes between HC and PD groups. **I-K.** Reporting of alterations in PBMC, myeloid cell-and T cell-mediated responses stratified by intervention type. Cell responses indicate the various cytokine outputs, including IFNγ, TNF, IL-17α, IL-1α, IL-1β, IL-2, IL-5, IL-6, IL-8, IL-10. The color indicates the directionality of changes in PD relative to HC. HS, human serum; FBS, fetal bovine serum; TCR, T-cell receptor; ELISA, enzyme-linked immunosorbent assay; PCR, polymerase chain reaction; ROS, reactive oxygen species.

Since LRRK2 is highly expressed in neutrophils, monocytes (Fan et al., 2018), but also B-cells (Thevenet et al., 2011), fifteen *in vivo* studies have utilized blood immune cells to study LRRK2-related changes. Nine of them evaluated LRRK2 kinase activity and explore LRRK2 as a biomarker (Fig. S2). The authors reported increased LRRK2 activity in genetic G2019S-LRRK2 PD, while data on LRRK2 activity or expression in sporadic PD was contradictory. Three articles studied LRRK2-related molecular pathways and another three investigated the role of LRRK2 in cellular processes such phagocytosis and endocytosis although they were not particularly focused on PwPD. (Fig. S2).

The adaptive immune compartment exhibited similarly heterogeneous yet some reproducible patterns of dysregulation across studies, with findings spanning both cell proportions and molecular features (Fig. 4C, see Table 2B for the list of included studies). In terms of cell proportions, overall lymphocytes frequencies were reduced across studies, with T cells (CD3+) reported to be less abundant in PD (Fig. 4F). Fewer studies have described B-cell abundance in PD relative to T lymphocytes albeit such studies consistently documented a decrease in naïve B cells concomitant with an increase in specific subsets of regulatory (CD5+CD19+), double-negative (CD27-IgD-), non-switched memory (CD27+IgM+IgD+) and activated (TNF+CD19+) B cells. By contrast, T-cell populations showed a mixed profile, with changes in cell proportions varying depending on the specific subtypes being assessed, and even across studies. Despite the heterogeneity in T lymphocyte findings, 11 studies described a higher abundance of cytotoxic or pro-inflammatory effector CD8 and/or CD4 subsets in PD, whereas the data examining naïve T-cell populations is more contradictory between studies. These emerging patterns highlight a recurring theme across studies – an expansion of activated lymphocyte populations. Potential skewing toward cytotoxic/pro-inflammatory T-cell programs further aligns with various reports on molecular features indicating elevated expression of IL-6, TNF, IFN-γ in PD lymphocytes (Fig. 4G; in light purple). However, several transcription factors central to T-cell differentiation (TBX21, STAT1/3/4/6) showed more nuanced results with very few papers interrogating these proteins. Furthermore, dopamine-related markers, including DRD3 and DAT, also differed between PD and matched controls across studies (Fig. 4G; in pink), suggesting an impaired dopaminergic signaling in circulating lymphocytes. Furthermore, increased frequency of DAT+ PBMCs is associated with worsening motor score indicative of disease relevance (Gopinath et al., 2024). Other aberrant adaptive immune *states* include markers of oxidative stress, which are consistently upregulated in PD and analogous to observations in innate immune studies (Fig. 4E, G; in green).

Together, these observations underscore substantial methodological heterogeneity, but also reveal convergent immune signatures, marked by altered subset frequencies and shifts in molecular features that suggest a primed inflammatory activity across both innate and adaptive immune compartments in PD.

### Overview of ex vivo studies illustrating peripheral immune traits in PD

To further contextualize immune findings, we examined methodological characteristics of studies employing *ex vivo* assays to interrogate peripheral immune cell functions in PD (see Table 4 for the list of included studies). These experiments involve isolating PBMCs from patients and healthy donors, followed by culture and stimulation under defined conditions, and then application of a plethora of immune phenotyping approaches (Fig. 5A). Sample handling practices varied, with most studies using freshly processed cells rather than cryopreserved samples (Fig. 5B). Culture conditions were inconsistently described; approximately 60% used RPMI supplemented with fetal bovine serum (FBS), a smaller proportion employed human serum (HS) or other formulations, and roughly 20% did not specify media composition (Fig. 5C). Stimulation protocols also differed, with the majority applying 1-3 days of exposure, about 30% performing acute stimulations of less than 24 hours, and only a few extending beyond 3 days (Fig. 5D). Most studies analyzed a single timepoint rather than multiple sampling intervals (Fig. 5E), limiting insight into dynamic immune responses. In some studies, specific populations are either positively or negatively selected and then plated *ex vivo* in comparison to plating bulk PBMCs in which the presence of multiple cell types is likely to influence the phenotype of specific subsets of cells being studied.

A wide range of stimuli were employed, with toll-like receptor (TLR) agonists and other target-specific pharmacological agents (such as NFAT, ATP and NLRP3 inhibitors, as well as inflammasome and redox state agonists) being the most common interventions, followed by co-culture systems, T-cell receptor (TCR) agonists and synthetic peptide pools. Fewer than ten functional studies applied genetic manipulations, CD40 agonists, synthetic DNA, viruses, bacteria or cytokines (Fig. 5F). Correspondingly, a variety of analytical techniques have been utilized to evaluate functional outcomes. Flow cytometry and ELISA dominated methodological approaches, while multiplex platforms, qPCR, Western blotting, and high-throughput sequencing were less commonly performed (Fig. 5G). Functional readouts primarily centered on general cytokine production by PBMCs, followed by assays probing T cell-and myeloid cell-mediated responses (Fig. 5H). Other measures included assessments of cell proliferation, cell death, mitochondrial function, reactive oxygen species (ROS) production and B-cell mediated responses. Further stratification of functional readouts by intervention type revealed that both total PBMC and myeloid cell-specific cytokine production were most frequently examined following TLR agonists challenge (Fig. 5I, J), whereas T cell-oriented studies favored peptide pools stimulation to evaluate antigenic responses (Fig. 5K).

Notably, T-cell hyperresponsiveness is the most reproducible immune *trait* in PD (Fig. 5K), whereas myeloid and bulk PBMC cytokine outputs show more variable patterns across multiple interventions – ranging from increased, decreased, unchanged or bidirectional changes in PD versus HC (Fig. 5J). Stimulation-based assays implicating CD4+ and/or CD8+ T cells revealed exaggerated cytokine production, with repeated reports of elevated secretion of TNF, IFN-γ, IL-17A, IL-5 and IL-10 in PD lymphocytes following treatment with mitogens (such as phytohemagglutinin, PHA), α-synuclein/viral peptides, Th1-polarizing cytokines, and after JKAP knockdown – a protein tyrosine phosphatase involved in suppressing T-cell immunity. Accordingly, JKAP loss induced cell surface expression of early activation markers CD25 and CD69 in circulating CD4+ T cells from PwPD. While monocytes exhibit both heightened and blunted responses depending on the stimulus, among all cytokines/molecules reported (IL-1β, IL-1α, IL-8, IL-10, TNF and soluble-CD163), IL-6 production was often measured across all *ex vivo* myeloid cell studies, noting its relevance when assessing innate immunity. Upon comparing interventions, extracellular vesicles stimulation promoted enhanced PD myeloid cell activation in at least two studies, whereas reduced cytokine output occurred in response to mitogens (such as concanavalin A) and TLR2 agonist (Pam3Cys). Notably, both lipopolysaccharide and α-synuclein challenges induced distinct myeloid cell responses across different studies, resulting in either decreased or increased in PD *versus* HC. These bidirectional changes emphasize that the participation of innate immunity in PD is highly context-and ligand-dependent, possibly reflecting differential TLR engagement/expression or altered distribution of classical, intermediate and non-classical subsets which are often not defined in such functional assays. Moreover, bioenergetic changes can also account for the differences as such process are energetically demanding.

Overall, these findings demonstrate reproducible and robust differences between control and PD PBMCs despite the inherent limitations of methodological differences in cell isolation and handling, *ex vivo* culture conditions, stimulation paradigms, and analytical techniques across studies, which undoubtedly influence and contribute to the reported functional heterogeneity in immune phenotypes in PD.

## Discussion

Across nearly five decades of research, our scoping review underscores the emerging consensus that peripheral immune populations exhibit evidence of dysregulation in PD, with robust differences detectable across multiple sites and numerous assays, although the reported magnitude, direction, and cellular specificity of these alterations vary widely. Our synthesis postulates that this variability is not solely biological but could be amplified by differences in cohort composition, clinical characterization, blood sample handling, anticoagulant selection, cell state, and analytical platforms. Inadequate reporting of critical parameters, including centrifugation conditions, flow cytometry gating strategies, culture media formulations, and viability measures further limits reproducibility and complicates meta-level interpretation. Of note, despite the implementation of a rigorous and systematic search with carefully constructed keywords, a handful of relevant studies containing additional information exist but were not captured within the databases and search parameters used. To this end, we have manually added such studies curated by experts in the field. This nonetheless highlights a potential gap in the Findable aspect of FAIR guiding principle (Wilkinson et al., 2016), indicating that despite being present, some research outputs remain difficult to locate.

Reproducible changes in NLR levels identified from studies employing *rudimentary immunophenotyping* are among the notable consistent *in vivo* immune signatures described in this review. It is however noteworthy to take caution when interpreting NLR data in PD as it is commonly used as diagnostic markers for inflammation. The sensitivity of immune cells to diverse environmental factors (such as infections) highlights a key challenge when characterizing the role of peripheral immunity in PD. Yet, emerging studies reveal that presenting an immune challenge to PBMCs or enriched populations from peripheral blood uncovers dysfunction in immune responsiveness which may contribute to the heterogeneity of PD endophenotypes (Hartlova et al., 2018; Mark, Staley, et al., 2025; Mark, Titus, et al., 2025; Wallings et al., 2024).

Our overview of results from *in vivo* studies unifies key patterns suggestive of an immune system in a “heightened alert” state in PD (Fig. 6). These include increased frequencies and activation states of cytotoxic and pro-inflammatory T-cell subsets, altered monocyte subset distributions, and changes in molecular markers of antigen presentation, T-cell engagement, chemotaxis, and innate inflammatory signaling. However, distinguishing disease-related stable immune features from transient *states*, that may fluctuate with environmental exposures, stress, circadian rhythms, or medication, can be difficult considering the predominance of single-timepoint sampling. This challenge is magnified by the fact that many PD cohorts include individuals on dopaminergic therapies, whereas healthy donors do not. Since dopamine modulates immune signaling (Feng & Lu, 2021), comparing treated PD patients to untreated controls introduces ambiguity as to whether observed differences, particularly in proteins involved in dopamine transport and/or synthesis, reflect disease mechanisms, treatment effects, or both.

**Figure 6:**
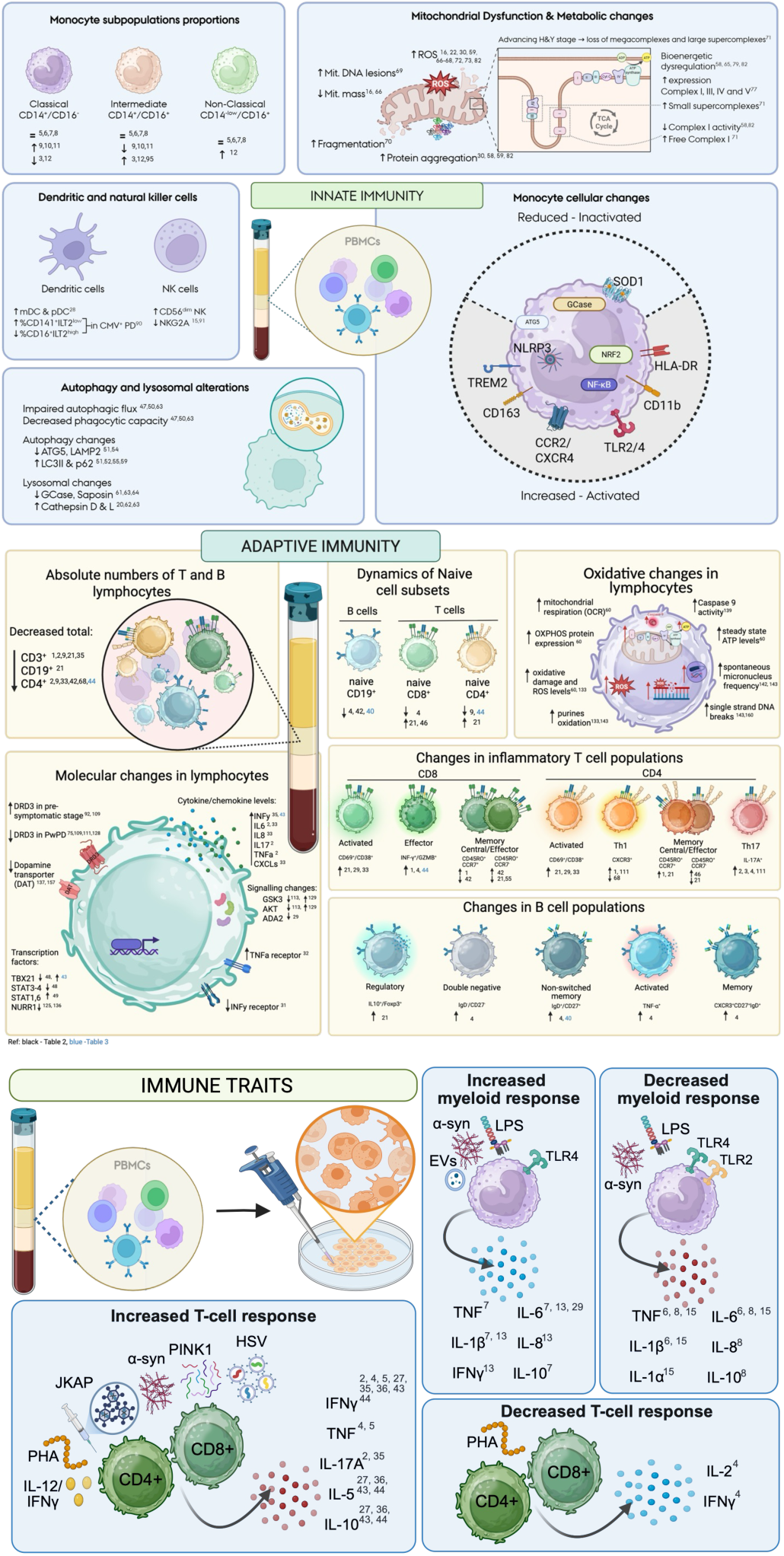
Synthesis of peripheral immune alterations in Parkinson’s disease (PD). Graphical summary of key findings from *in vivo* and *ex vivo* studies of peripheral blood immune cells in PD. Innate immune changes include altered monocyte subset distribution, mitochondrial and metabolic dysfunction, impaired autophagy-lysosomal pathways, and dysregulated inflammatory signaling. Adaptive immunity is characterized by reduced circulating lymphocyte numbers, shifts in naïve and memory T-and B-cell subsets, and enhanced pro-inflammatory polarization. Functional assays reveal heterogenous myeloid and T-cell responses to inflammatory stimuli. Together, these findings support systemic immune dysregulation in PD involving both activation and functional impairment.

To unveil facets of immune perturbations in PD that may be difficult to parse out at baseline, we summarized *ex vivo* studies which offer additional insight into more stable immune phenotypes (Fig. 6). A key advantage of these studies is that stimulus-evoked activation and resolution responses, defined as immune *traits*, can be elicited in a controlled experiment regardless of aforementioned exogenous factors. Indeed, we found that a variety of stimuli similarly induced T-cell hyperresponsiveness in PD, while myeloid cell responses are highly stimulus-dependent, with cytokine release increasing or decreasing in PD depending on the ligand, duration of stimulation, technique utilized, and monocyte subsets present in the culture. Additionally, stimulation-based assays have the potential to increase energetic demand, that may unleash deficits in immune cell bioenergetics, which have been strongly implicated in PD pathogenesis (Mark & Tansey, 2025). This was noted in our review with some *ex vivo* studies reporting metabolic alterations involving cell proliferation, apoptosis, mitochondrial functions and oxidative stress. Consistent with some of these phenotypes, findings from several *in vivo* studies indicate that mitochondria in PBMCs from PwPD may be hyperactive but inefficient, potentially contributing to a toxic cellular environment and chronic inflammation.

Evidence for a hindered autophagy-lysosomal clearance system in PD peripheral immune cells was striking and may point toward compromised intracellular vesicle dynamics. If vesicular trafficking is largely impaired, immune cells could exhibit an inflamed basal transcriptional state while being less capable of secreting cytokines, depending on the assay readout. This model reconciles apparent discrepancies across studies: intracellular cytokine staining using flow cytometry-based methods may show elevated levels, whereas secretion-based assays (such as ELISA) may show blunted cytokine release. Differences in experimental approach could therefore partly underlie the variability in reported functional outcomes. Moreover, alterations in immune activity, vesicular machinery, degradation systems and metabolism are also mirrored through transcriptomics analyses. However, such studies, particularly single-cell omics approaches, remain relatively scarce and underpowered, emphasizing the need for larger, harmonized datasets.

Intriguingly, certain molecular features, such as defective autolysosomes, are shared between idiopathic and genetic forms of the disease as exemplified by GBA mutant carriers showing reduced expression in PBMCs of key lysosomal proteins implicated in autophagy (Saposin C and LAMP2) (Avenali et al., 2021; Papagiannakis et al., 2015). Similarly, bulk RNAseq studies detected dysregulated pathways involving mitochondrial function and vesicle trafficking in both idiopathic PD and LRRK2 mutation carriers (Bolen et al., 2025; Mutez et al., 2014). Nevertheless, distinct immune signatures involving glucocorticoid receptor and lymphocyte signaling pathways were also identified between these two PD subtypes (Mutez et al., 2014). LRRK2 kinase activity in peripheral immune cells provides additional granularity. LRRK2 G2019S carriers have heightened PBMC kinase activity but not in idiopathic PD, while sex-dimorphism associated with LRRK2 suggest potential gene-sex interactions shaping immune phenotypes (Melachroinou et al., 2020). A more comprehensive synthesis of subtype-specific immune signatures in PD is thus warranted to understand potential shared and distinct pathogenic mechanisms across genetic backgrounds.

Another major future direction will require systematically resolving how immune changes differ across disease severity. In this review, we listed the proportions of studies enrolling participants at distinct PD stages and noted most have recruited moderate-to-late stage patients, defined by disease duration >5 years and/or clinical measures attributed to low motor performance (H&Y stages III-V, Webster ratings >8, Braak stages III-V, UPDRS-III ratings >18), whereas only a handful have profiled immune changes at the prodrome. Several immune features described here vary with disease progression, such as monocyte frequencies and myeloid cell surface markers (*e.g.,* CD11b, CCR2, TLR2/4) (Konstantin Nissen et al., 2022; Thome et al., 2021; Wijeyekoon et al., 2020). Many of these abnormalities appear strongest in early, untreated PD and diminish with dopaminergic therapy (Ciaramella et al., 2013; Solini et al., 2021a), raising questions about whether treatment and/or disease severity modulates immune signaling or whether chronic disease leads to immune exhaustion. Meanwhile, other receptors are reported to increase with time as the disease progresses, such as HLA-DR and soluble CD163, although it is unclear if their signaling efficiency remains at later stages (Konstantin Nissen et al., 2022; Thome et al., 2021; Wijeyekoon et al., 2020). Longitudinal studies in treatment-naïve cohorts or prodromal stage are essential to disentangle these possibilities.

Taken together, the literature paints a picture of a peripheral immune system that is activated, imbalanced and metabolically stressed in PD, particularly early in the disease course. However, these changes remain obscured by methodological artifacts and by challenges that impede biologically meaningful interpretation. To build reliable immune endophenotypes in PD and guide the development of immune-modulating therapies, a shift is needed toward studies that (i) adopt standardized reporting (ii) implement unified sample processing workflows, (iii) integrate multimodal immune profiling across diverse and well-characterized cohorts, (iv) interrogate cell type-specific features rather than bulk measures, and (v) distinguish stable immune *traits* from transient immune *states*. Future integrative reviews comparing across genetic forms and stages of PD will help further resolve disease heterogeneity.

## Illustrations

Graphical representations of *in vivo* and *ex vivo* studies were created with BioRender.

## Author contributions

SJR and LL performed initial screening of the literature. SJR, JJP, NL, ZV and MOO extracted information in all included studies. SJR drafted the manuscript and prepared figures. JAS, MGT, MRR, KE and AH provided funding, supervised the work, and revised the final manuscript. All authors contributed to the article and approved submitted version.

## Data availability

Raw tabular data is available at https://zenodo.org.

---

**Supplementary Figure 1:**
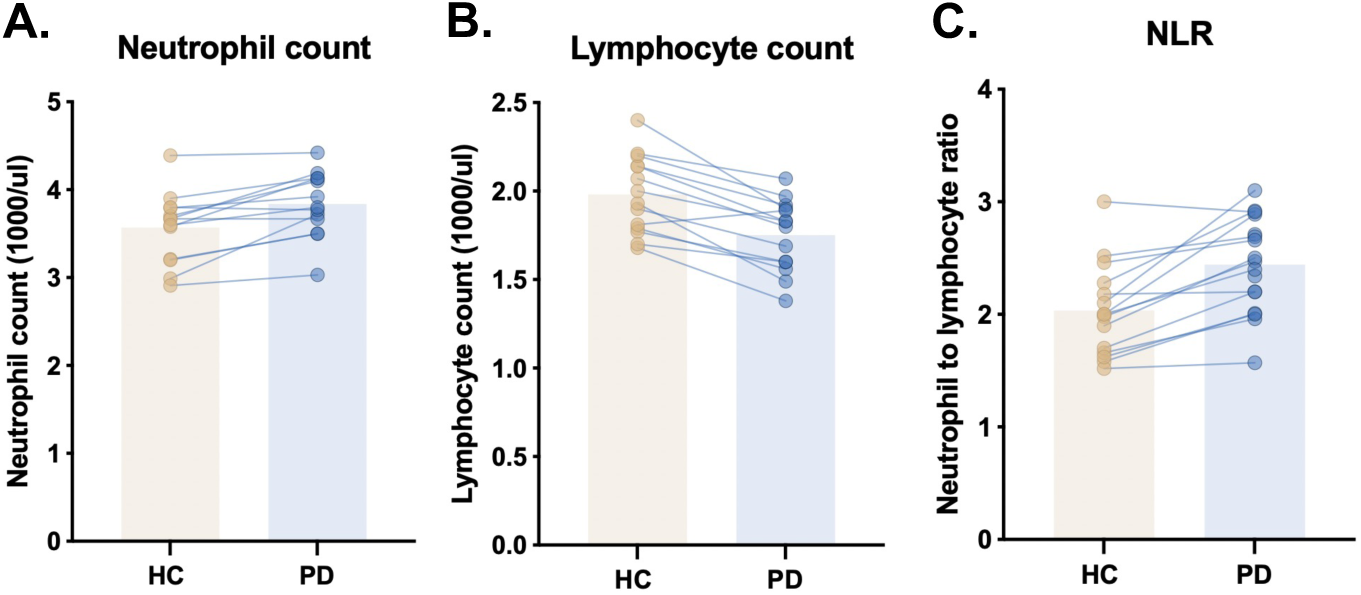
Elevated peripheral blood neutrophil-to-lymphocyte ratio in Parkinson’s disease revealed by rudimentary immunophenotyping. A-C. Neutrophil and lymphocyte cell counts, as well as neutrophil-to-lymphocyte ratio (NLR) detected in peripheral blood of healthy control (HC) and PD patients. Paired data points represent comparisons in each study.

**Supplementary Figure 2:**
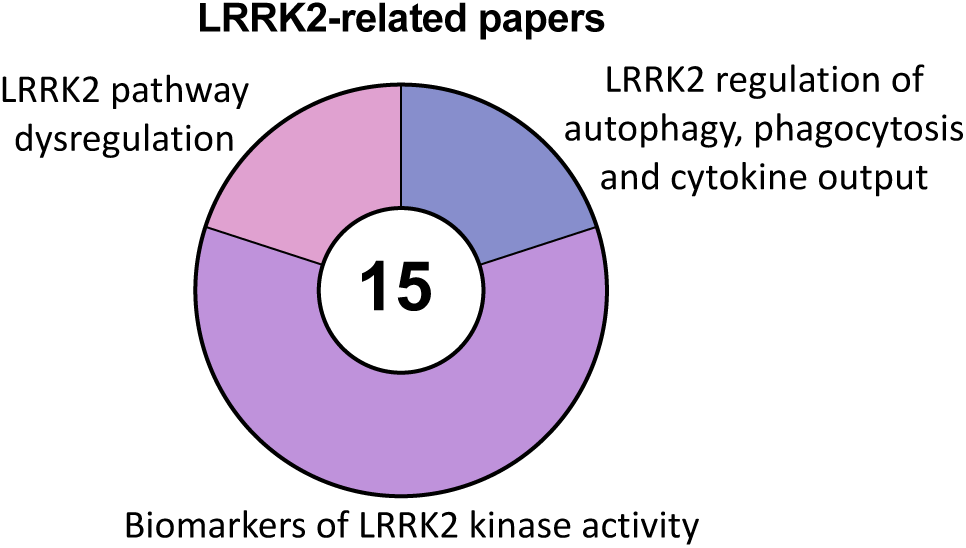
Synthesis of LRRK2-related changes in Parkinson’s disease peripheral immune cells. A pie chart depicting the number of papers employing peripheral blood mononuclear cells to characterize LRRK2 kinase activity, LRRK2-related molecular pathways, and LRRK2’s role in cellular processes, such as autophagy, phagocytosis and cytokine output.

**Supplementary Table 1:** Sample database strategy for PubMed (24-02-2025)

| Search Concept | Search Terms |
| --- | --- |
| PD/RBD/LBD (1) | ("Parkinson's Disease" OR "Parkinson Disease" OR "Parkinsonism" OR "REM Sleep Behavior Disorder" OR "Sleep Paralysis" OR "REM Sleep Parasomnias" OR "Lewy Body" OR "Synucleinopathies" OR "Synucleinopathies" OR "alpha Synuclein Pathology") |
| Sources (2) | ("CSF" OR "Cerebrospinal Fluid" OR "Blood" OR "Serum" OR "Plasma" OR "Feces" OR "Stool" OR "Saliva" OR "Skin" OR "Gut" OR "Intestine" OR "Gastrointestinal Tract" OR "Ileum" OR "Cecum" OR "Anal" OR "Colon" OR "Rectum" OR "Jejunum") |
| Inflammation (3) | ("Inflammation" OR "Immune response" OR "Immune process" OR "Immunity" OR "Adaptive immunity" OR "Innate immunity" OR "Antigen-mediated" OR "Humoral response" OR "Cellular response" OR "Autoimmunity" OR "Cross-presentation" OR "Trained Immunity" OR "Mucosal immunity" OR "T-Cell Antigen Receptor" OR "Cytokines" OR "Chemokines" OR "Macrophage Inflammatory Proteins" OR "Growth Differentiation Factor 15" OR "Hematopoietic Cell Growth Factors" OR "Colony-Stimulating Factors" OR "Hepatocyte Growth Factor" OR "Interferon" OR "Interleukin 1 Receptor Antagonist Protein" OR "Interleukins" OR "Leukemia Inhibitory Factor" OR "Lymphokines" OR "Monokines" OR "Oncostatin M" OR "Osteopontin" OR "Thymic Stromal Lymphopoietin" OR "Transforming Growth Factor beta" OR "Tumor Necrosis Factor" OR "4-1BB Ligand" OR "B-Cell Activating Factor" OR "Ligand" OR "CD27" OR "CD30" OR "CD40" OR "FAS" OR "OX40" OR "RANK" OR "Lymphotoxin" OR "Exhaustion" OR "Cytokine signaling" OR "Toll-like receptors" OR "TLR" OR "Pattern recognition receptor" OR "PRR" OR "Platelets" OR "Erythrocytes" OR "Hemocytes" OR "Leukocytes" OR "CD8" OR "CD19" OR "CD138" OR "CD14" OR "CD16" OR "T helper" OR "T cell" OR "Tcell" OR "T-cell" OR "TH1" OR "Th1" OR "TH17" OR "Th17" OR "TH2" OR "Th2" OR "Treg" OR "Monocyte" OR "Peripheral blood mononuclear cells" OR "PBMC" OR "Lymphocytes" OR "Macrophage" OR "Dendritic cells" OR "Neutrophil" OR "Natural killer" OR "Myeloid cells" OR "Phagocytes" OR "M0" OR "M1" OR "M2" OR "Mreg" OR "Antigen-presenting cells" OR "APC") |
| Combined (4) | (1) AND (2) AND (3) |

